# Multifunctional PolySpermine-based Nanocapsules for Targeted Gene Delivery to Gastric Cancer Cells

**DOI:** 10.1101/2025.08.27.672643

**Authors:** Sahar Mohajeri, Mahsa Raei, Shima Bourang, Mehran Noruzpour, Hashem Yaghoubi

**Affiliations:** Department of Chemistry, Ard.C., Islamic Azad University, Ardabil, Iran; Department of Biology, Ard.C., Islamic Azad University, Ardabil, Iran; Department of Agronomy and Plant Breeding, Faculty of Agriculture and Natural Resources, University of Mohaghegh Ardabili, Ardabil, Iran

**Keywords:** Gene delivery systems, Nanocapsules, Polycations, Gastric neoplasms, Transfection, Drug carriers

## Abstract

In this study, multifunctional nanocapsules were developed and evaluated for targeted gene delivery to AGS gastric cancer cells. The design of the nanoparticles utilized hyperbranched polyspermine (HS) for efficient DNA condensation, polyethylene glycol (PEG) to increase nanoparticle stability and prolong circulation time via stealth properties, and dual-targeting ligands, i.e., folic acid and glucose, to improve selective binding and internalization by cancer cells. Folic acid targets folate receptors (FRα), while glucose binds glucose transporters (GLUTs), both of which are overexpressed in gastric cancer cells, thereby increasing uptake specificity. The synthesized ternary copolymers composed of polyspermine, PEG, folic acid, and glucose (PSPFG) were comprehensively characterized via multiple analytical techniques, including proton nuclear magnetic resonance (¹H-NMR), Fourier transform infrared spectroscopy (FTIR), thermogravimetric analysis (TGA), and derivative thermogravimetric (DTG) analysis, to confirm their chemical structure and thermal stability. After complexation with DNA, the PSPFG/100 DNA nanocapsules were analyzed by scanning electron microscopy (SEM) and transmission electron microscopy (TEM), which revealed uniform spherical nanoparticles with a nanoscale size. Dynamic light scattering (DLS) measurements confirmed a narrow size distribution, with an average particle size of 265 ± 18 nm. Biocompatibility assays using the 3-(4,5-dimethylthiazol-2-yl)-2,5-diphenyltetrazolium bromide (MTT) assay demonstrated significantly reduced cytotoxicity compared with the commonly used polyethylenimine (PEI) vector. Agarose gel electrophoresis revealed strong DNA binding, effective charge neutralization, and resistance to enzymatic degradation. Importantly, fluorescence microscopy and flow cytometry analyses demonstrated high transfection efficiency in AGS cells, with the optimized PSPFG50/DNA formulation achieving a transfection rate of 53.37%. These results collectively indicate that PSPFG-based nanocarriers exhibit favorable biocompatibility and enhanced gene delivery performance, addressing major limitations of traditional polycationic vectors. These findings suggest promising potential for the clinical translation of these spermine-derived nanocapsules in gastric cancer gene therapy.

## 1. Introduction

Gastric cancer, also known as stomach cancer, remains one of the leading contributors to cancer-associated mortality worldwide [1]. This malignancy often presents at an advanced stage due to its asymptomatic nature in early development, rendering timely diagnosis challenging [2]. The conventional methods for treating cancer include radiation therapy, chemotherapy, surgery, and targeted therapies [3]. However, these methods frequently have drawbacks because of their systemic toxicity, drug resistance, and unfavorable side effects, which calls for the investigation of more creative treatment approaches [4]. The AGS cell line, derived from human gastric adenocarcinoma, is widely utilized in research to study gastric cancer biology and evaluate new therapeutic agents [5]. This cell line serves as a valuable model for understanding the molecular mechanisms underlying gastric cancer progression and assessing the efficacy of novel therapies. Owing to their close resemblance to gastric tumors, AGS cells provide an essential platform for investigating potential treatments and drug delivery systems.

In recent years, nanotechnology has emerged as a transformative tool in oncology, particularly for enhancing the targeting and delivery of therapeutics [6]. Engineered nanocapsules for drug and gene delivery increase the solubility and bioavailability of therapeutic compounds while allowing for site-specific controlled release, which reduces systemic side effects and increases overall treatment effectiveness [7]. The targeted modification of nanocapsules for gene and drug delivery is a promising method that researchers use to overcome the weaknesses of conventional medical treatments [8]. The use of gene therapy provides new opportunities to treat genetic diseases and cancers previously deemed incurable [9]. Numerous biomolecular tools, including antisense oligonucleotides, small interfering RNAs (siRNAs) [10], and other nucleic acid-based agents, have been employed to modulate gene expression. Despite these efforts, the successful and efficient delivery of such molecules into mammalian tissues remains a considerable challenge. The large molecular weight and inherent negative charge of DNA significantly limit its capacity to cross cellular membranes [11]. These negatively charged nucleic acids can be complexed with cationic polymers, leading to the formation of nanocomposites with improved delivery efficiency [12].

Among the various biomaterials that have been explored and studied for their potential in gene delivery applications, nanoparticles made from polyamines have attracted considerable interest from researchers [13, 14]. This interest is largely due to their favorable biocompatibility, which allows them to function well within biological systems, and their inherent ability to condense nucleic acids effectively [15]. Additionally, polyamine-based nanoparticles have shown great potential for enhancing the cellular uptake of the genetic material they carry [16]. Spermine, a polyamine that is naturally present in living organisms, stands out for its favorable properties, which contribute to the stabilization and protection of nucleic acids during delivery [17]. Furthermore, the interactions of spermine with various cellular components can significantly enhance the intracellular delivery of therapeutic genes, thereby contributing to improved therapeutic outcomes in gene therapy approaches [18].

Another copolymer employed in gene delivery is polyethylene glycol (PEG), a hydrophilic polymer that has gained significant attention in the field of drug delivery, particularly for gene therapy and cancer treatment [19]. Owing to its unique physicochemical properties, including biocompatibility, low immunogenicity, and ability to increase the solubility of various therapeutic agents, PEG is an ideal candidate for formulating drug delivery systems, especially nanoparticles [20]. One of the most notable advantages of PEG is its “stealth” effect, which helps to evade detection by the immune system, thereby prolonging the circulation time of the drug in the bloodstream and improving its bioavailability [21]. In the context of gene delivery for cancer treatment, PEG is often used to modify the surfaces of nanoparticles, liposomes, or other delivery vehicles [5]. Modification of the nanoparticle surface significantly improves stability and decreases aggregation within biological fluids, facilitating more effective targeting of tissues [22]. Moreover, PEGylation can increase the permeability of nanoparticles through biological barriers, facilitating their uptake by cancer cells. The use of PEG in gene delivery systems also allows the incorporation of targeting ligands that can interact specifically with receptors overexpressed on cancer cells [23].

On the other hand, glucose has emerged as a promising component in gene delivery systems, particularly for enhancing the targeting and uptake of therapeutic genes by cancer cells [24]. Owing to the high metabolic demands of many cancer cells, which often exhibit elevated glucose consumption (a phenomenon known as the Warburg effect), incorporating glucose into gene delivery vehicles can exploit this metabolic preference [25]. By functionalizing nanocapsules or liposomes with glucose moieties, researchers can create targeted delivery systems that specifically bind to glucose transporters that are overexpressed on the surface of cancer cells. This strategy not only increases the efficiency of gene uptake but also minimizes off-target effects on healthy tissues. Furthermore, glucose-modified carriers can increase the stability and solubility of nucleic acids, facilitating their delivery into cells [26]. Folic acid (FA) and glucose were used in this study for secondary cellular receptors because of their improved biocompatibility, cellular uptake, targeting mechanism, and synergistic effects. In gene therapy and cancer treatment, combining the targeting properties of glucose and folic acid in gene delivery systems is a promising approach.

This research aimed to design multifunctional nanocapsules that enhance gene delivery efficiency to cancerous tissues while reducing off-target effects on normal cells. For this purpose, hyperbranched spermine (HS) was synthesized to improve the DNA-binding and condensation ability of native spermine. To promote prolonged systemic circulation and selective delivery to breast cancer cells, the branched polymer was subsequently modified with polyethylene glycol (PEG) conjugates of folic acid and glucose.

## 2. Materials and methods

### 2.1. Materials

Potassium chloride (KCl), dimethyl sulfoxide (DMSO), 3-(4,5-dimethyl-2-thiazolyl)-2,5-diphenyl-2H-tetrazolium bromide (MTT), ethanol, HCL, NaCl, KH_2_PO_4_, chloroform, and sodium hydroxide (NaOH) were purchased from Merck (Germany). Paraformaldehyde was prepared from Dr. Mojal Chemical Laboratory (Iran). Fetal bovine serum (FBS), RPMI 1640, and trypsin-EDTA were purchased from Biowest (USA).

### 2.2. Methods

#### 2.2.1. Synthesis of PolySpermine

For this purpose, first, 1 mmol of pentane-1,3,5-tricarboxylic acid was activated with EDC/NHS in dry DMF and converted to an NHS triester. Then, 3 mmol of spermine (with four amine groups) was added to the solution, and its amine groups reacted with NHS esters to form branched amide bonds. The reaction was carried out at 25-40°C under nitrogen gas. After completion of this step, the product was purified by precipitation in ether/hexane (**Figure 1**). The final product was collected and dried via a freeze‒drying process (Christ Alpha 1-4 LD plus, Germany).

**Figure 1.**
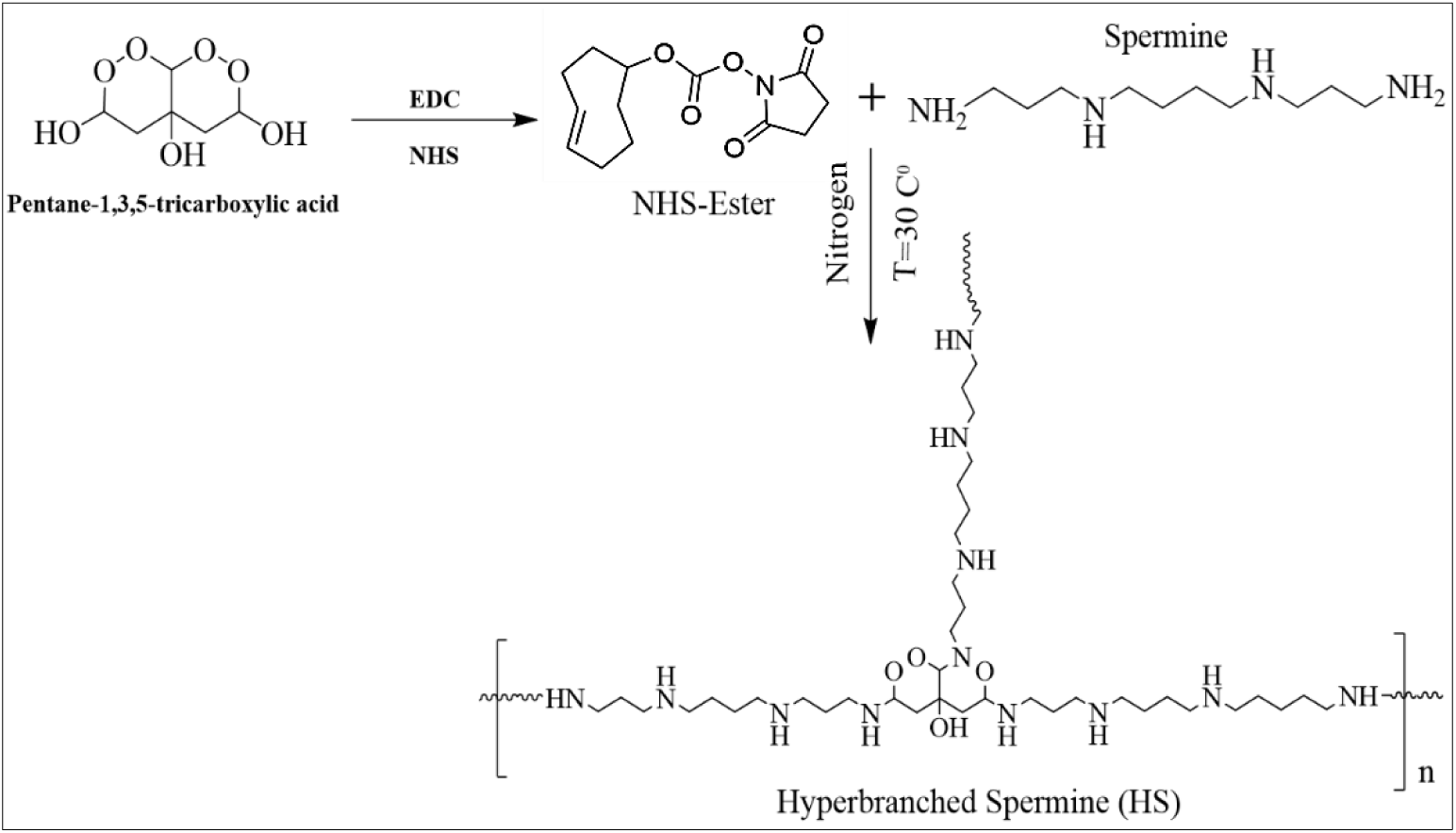
Schematic illustration of the synthesis of the HS.

#### 2.2.2. Synthesis of the PolySpermine-PEG-FA (PSPF) and PolySpermine-PEG-FA-Glu (PSPFG) nanocapsules

The PSPF copolymer was synthesized by reacting 1 g each of NH_2_-PEG-FA and NH_2_-hyperbranched spermine (NHS) with 50 mg of EDC and 40 mg of NHS in 10 mL of DMSO, followed by incubation at room temperature for 5 h. The reaction was continued for an additional 48 h after activated PEG-FA was added to the HS. After three days, the products were dialyzed against DMSO/water (**Figure 2**) [27].

**Figure 2.**
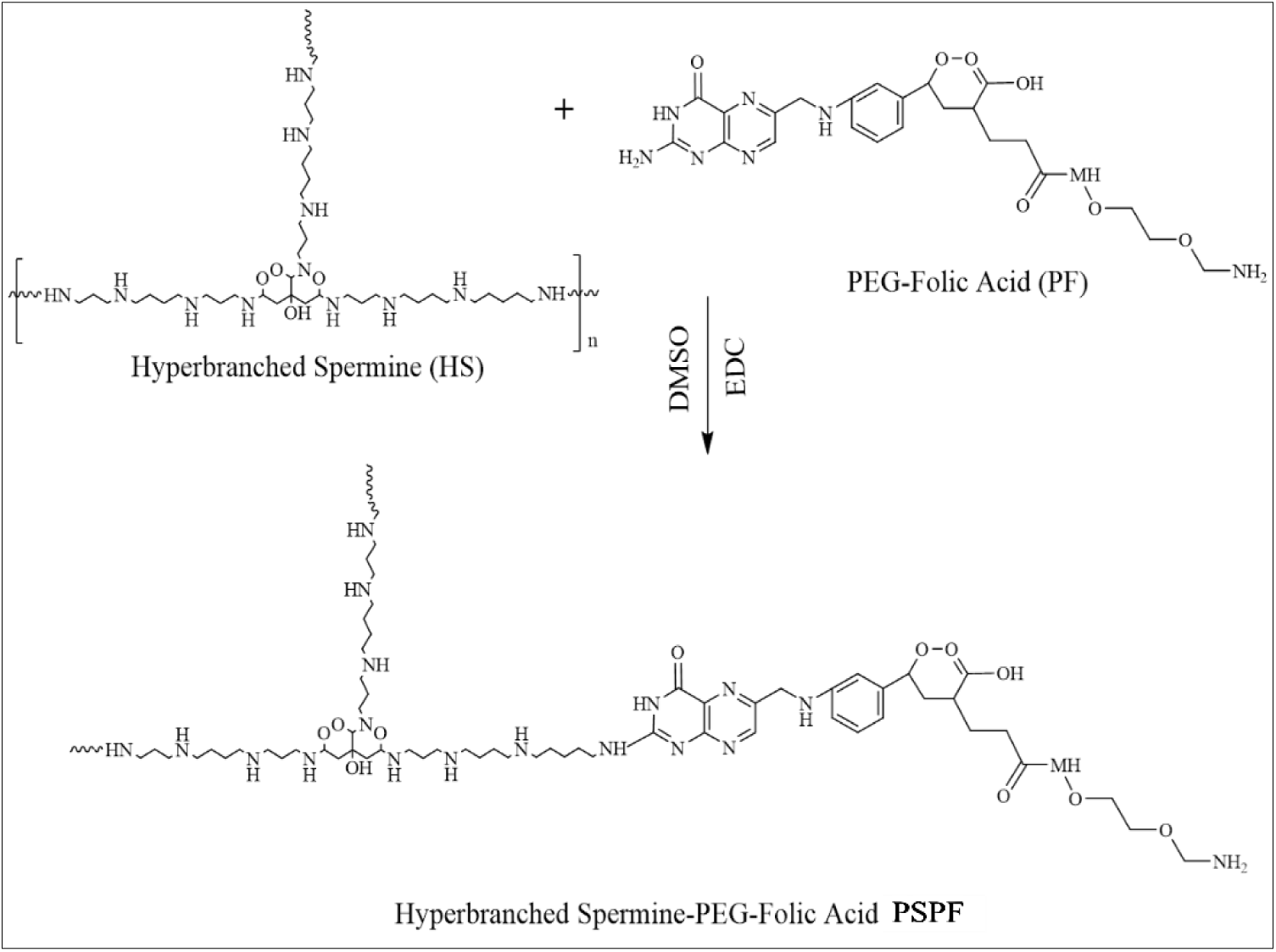
Schematic illustration of the synthesis of PSPF nanocapsules.

For the synthesis of PSPFG nanocapsules, the same PSPF polymer backbone was used (**Figure 3**).

**Figure 3.**
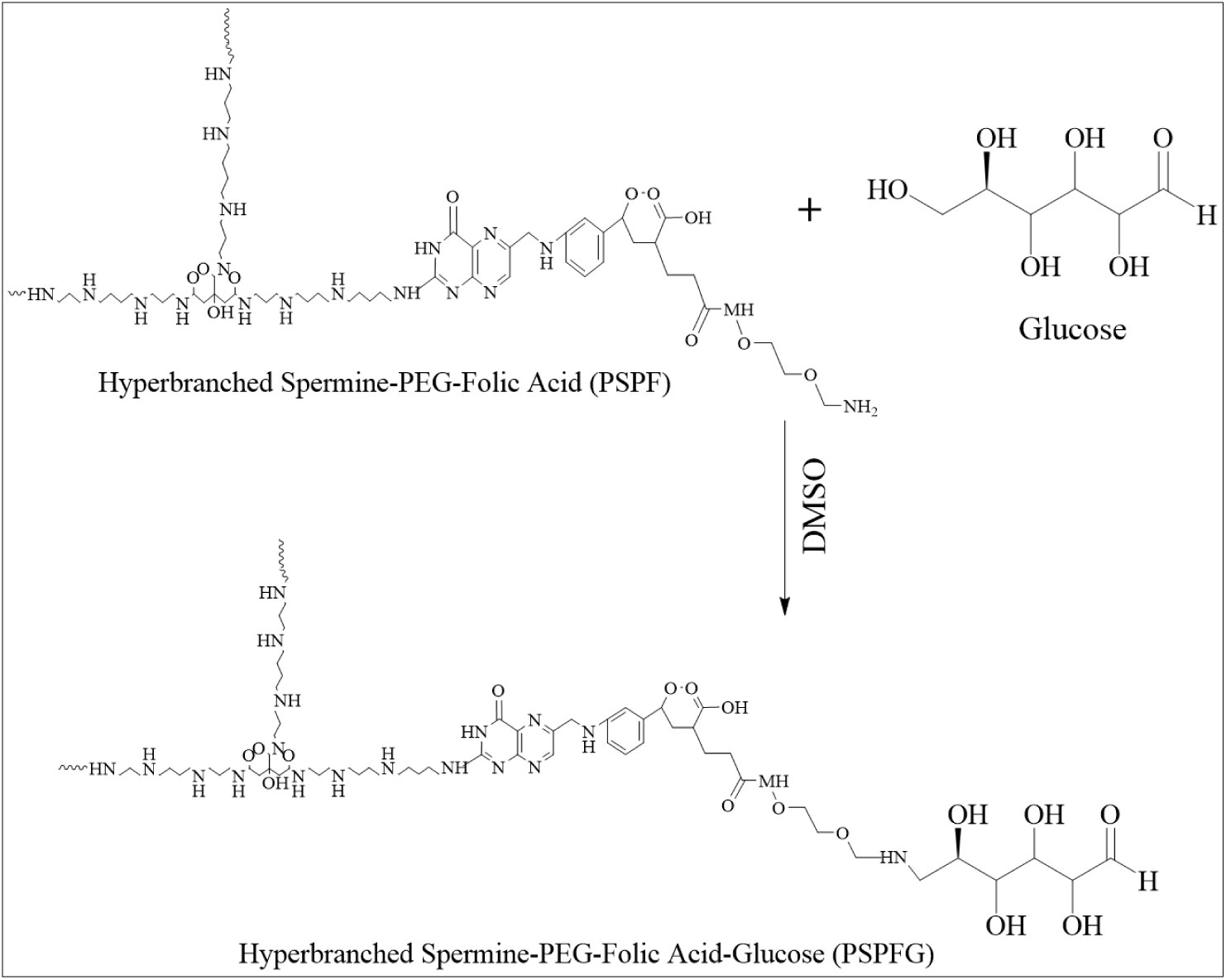
Schematic illustration of the synthesis of PSPFG nanocapsules.

#### 2.2.3. Preparation of PolySpermine-PEG-Folic acid/glucose (PSPFG)/DNA nanocapsules

PSPFG/DNA nanocapsules were prepared by mixing varying amounts of PSPFG with 5 μg of plasmid DNA in an appropriate buffer, followed by incubation at ambient temperature (∼25°C) for 30 min to allow self-assembly (**Table 1**) [28].

**Table 1.** Compounds used to prepare PSPFG/DNA nanocapsules.

| Nanocapsules name | PSPFG (µg) | DNA (µg) | PSPFG/DNA weight ratio (w/w/w %) |
| --- | --- | --- | --- |
| PSPFG5/DNA | 2.5 | 5 | (0.5:1) |
| PSPFG10/DNA | 5 | 5 | (1:1) |
| PSPFG25/DNA | 10 | 5 | (2:1) |
| PSPFG50/DNA | 25 | 5 | (5:1) |
| PSPFG100/DNA | 50 | 5 | (10:1) |
| PSPFG200/DNA | 100 | 5 | (20:1) |

#### 2.2.4. Characterization of nanocapsules

Infrared spectroscopy (FTIR-ABB Bomem-MB) and proton NMR spectroscopy (^1^H-NMR-Bruker 400 MHz) were used to analyze the produced copolymers. Transmission electron microscopy (TEM, JEM-2100) and dynamic light scattering (DLS, Malvern Instruments, Westborough, MA, USA) were used to analyze the nanocapsule shape, particle size, and zeta potential, respectively. Thermogravimetric analysis (TG-DTA-32, Japan) was used to investigate the thermal characteristics of spermine, polyspermine, and polyspermine-PEG-glucose at temperatures ranging from 25°C to 600°C with a heating rate of 20°C/min under atmospheric pressure [13].

#### 2.2.5. Determination of the DNA encapsulation efficiency of synthesized nanocapsules

The DNA encapsulation capability of various nanocapsules, including PSPG100/DNA, PSPF100/DNA, PSPFG100/DNA, and PEI100/DNA (the concentrations of each sample were prepared on the basis of the calculation path in **Table 1**. For example, for PEI100/DNA, the concentrations of DNA and PEI (1DNA:PEI 10) were considered) and were assessed via a spectrophotometer by measuring the absorbance of their suspension supernatants at 260 nm. The encapsulation efficiency was calculated by comparing the measured absorbance to the initial drug amount (1 mg) via the following formula [29]:

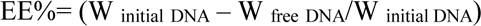

#### 2.2.6. Kinetics of the release of DNA from nanocapsules

The release profiles of DNA from PSPG100/DNA, PSPF100/DNA, PSPFG100/DNA, and PEI100/DNA nanocapsules were assessed under two distinct pH conditions: acidic (pH 5.5) and neutral (pH 7.4). Additionally, release studies were conducted in the presence of dextran sulfate at concentrations ranging from 0 to 100 μg/mL (0, 10, 20, 30, 40, 50, 60, 70, 80, 90, and 100 μg/mL).

For these experiments, PSPG100/DNA nanocapsules were initially incubated in 20 mL of phosphate-buffered saline (PBS) at 37°C. After centrifugation at 16602 × g for 30 min, the supernatant was collected for subsequent analysis, and the resulting nanocapsule pellet was gently resuspended in fresh buffer, followed by continued incubation under the designated conditions [19].

A Nanodrop spectrophotometer with a wavelength of 260 nm was used to measure the DNA concentration in each sample. The cumulative percentage of DNA released from the nanocapsules was calculated as follows:

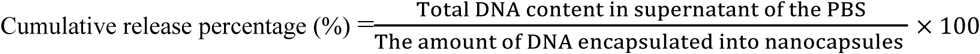

#### 2.2.7. Cell culture

The AGS human gastric cancer cell line (NCBI, C131) was sourced from the National Cell Bank of Iran, Pasteur Institute. The cells were cultured in RPMI 1640 medium (Gibco, USA) supplemented with 10% heat-inactivated fetal bovine serum (FBS) in a humidified incubator at 37°C with 5% CO_2_. Upon reaching 70–80% confluence, the cells were subcultured with trypsin-EDTA (Gibco, USA) to maintain exponential growth [5].

#### 2.2.8. *In vitro* cytotoxicity studies

The MTT assay is a commonly used technique for assessing cytotoxicity. It operates on the principle that viable cells convert 3-[4,5-dimethylthiazol-2-yl]-2,5-diphenyl tetrazolium bromide (MTT) into insoluble formazan crystals, reflecting their mitochondrial activity. Since mitochondrial function typically halts upon cell death, the MTT assay serves as a reliable indicator of cell viability [30].

In this study, the cytotoxic effects of PEI, PEI100/DNA (N/P=5), PSPG, PSPF, PSPFG, and PSPFG100/DNA nanocapsules were evaluated via the MTT assay. AGS cells were cultured in 96-well plates at a density of 7 × 10^3^ cells per well in 200 μL of complete RPMI 1640 medium supplemented with 10% fetal bovine serum (FBS). After a 24-hour incubation at 37°C in a humidified atmosphere with 5% CO_2_ to facilitate cell adherence, varying concentrations of nanocapsules (ranging from 0 to 100 µg/mL) were applied. The plates were then incubated for an additional 24 hours under identical conditions. Cell viability was evaluated via the MTT assay, and the absorbance was measured at 570 nm via a BioTek microplate reader (Winooski, VT, USA). The percentage of viable cells was determined according to the following formula [31]:

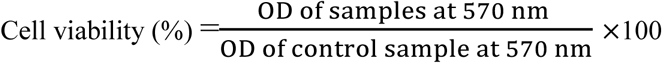

#### 2.2.9. Cell Apoptosis Analysis

Apoptosis involves the movement of phosphatidylserine from the inner to the outer plasma membrane, making it detectable with annexin-based fluorescence staining [32]. To distinguish apoptosis from necrosis, annexin is used in conjunction with propidium iodide.

In this study, AGS cells treated with PEI100/DNA and PSPFG100/DNA nanocapsules were analyzed for apoptosis via flow cytometry and an Annexin V-Dy634 kit (Immunostep) following the supplier’s protocol. Untreated cells were used as a positive control. The data were processed with a FACS Verse flow cytometer and FlowJoTM version 10 software [33].

#### 2.2.10. Agarose gel electrophoresis of nanocapsules

To determine the DNA-binding efficiency of PSPFG nanocapsules, agarose gel electrophoresis was carried out at various concentrations (5, 10, 25, 50, 100, and 200 µg). A 0.8% (w/v) agarose gel was prepared, and 2 µg of DNA, either encapsulated or free, was loaded per lane. Electrophoresis was run at 80 V for 60 minutes to visualize DNA mobility [34].

#### 2.2.11. DNA protection assay

The stability of DNA in PSPFG/DNA nanocapsules against degradation by plasma enzymes was tested by incubating 2 μg of DNA per sample, in either the nanocapsules or the free form, with 50 μL of 20% human plasma at 37°C for 30 minutes. Nuclease activity was inhibited via the addition of 5 μL of 0.5 M EDTA (pH 8), and heparin (1% w/v) was added before incubation for 4 hours at 37°C with shaking. The DNA was then examined via 2% agarose gel electrophoresis at 80 V [34].

#### 2.2.12. Transfection assay

Transfection serves as a key technique in molecular biology, allowing researchers to introduce exogenous nucleic acids into cells for functional analysis [35]. In this study, we compared the transfection efficiency of PEI/DNA (N/P 5:1) and PSPFG/DNA nanocapsules in AGS gastric cancer cells.

AGS cells (2 × 10^4^) were seeded into 24-well plates containing 1 mL of RPMI 1640 with 10% FBS and incubated at 37°C and 5% CO_2_ for 24 hours. The cells were then washed and transfected with 2 μg of DNA per nanocapsule in 1 mL of medium, either with or without serum. The controls included uncoated DNA and PEI/DNA complexes. After 7 hours, the medium was replaced, and the cells were incubated for an additional 48 hours. Flow cytometry (CyFlow Space, Germany) and inverted fluorescence microscopy (Nikon TE200) were used to assess pEGFP-N1 expression and overall transfection performance.

#### 2.2.13. Statistical analysis

Each measured parameter in this study was evaluated in triplicate or more. Data analysis was performed via one-way ANOVA via SPSS software, followed by Duncan’s test for multiple comparisons at a significance threshold of 0.05. The normal distribution of the data was examined via the Kolmogorov–Smirnov test. The results are reported as the mean values with standard deviations (means ± SD).

## 3. Results

### 3.1. ^1^H-NMR Results

The ^1^H-NMR spectrum of the PSPF copolymer revealed characteristic peaks at 1.7 and 2.1 ppm, corresponding to the methylene protons in the carbon backbone and the N-H protons of the polyspermine segments, respectively (**Figure 4 A**, peaks 1 and 2). Additional signals were detected at 2.8 and 3.7 ppm, attributed to the N–H proton and the hydroxyl protons in the main chain of PEG, respectively (**Figure 4 A**, peaks 3 and 4). The aromatic region between 6.0 and 8.0 ppm displayed peaks assigned to the protons of the folic acid moiety (**Figure 4 A**). Upon the conjugation of glucose to the PSPF copolymer, additional resonances appeared in the same 6.0–8.0 ppm range, which were attributed to the glucose-derived protons (**Figure 4 B**, peak 7).

**Figure 4.**
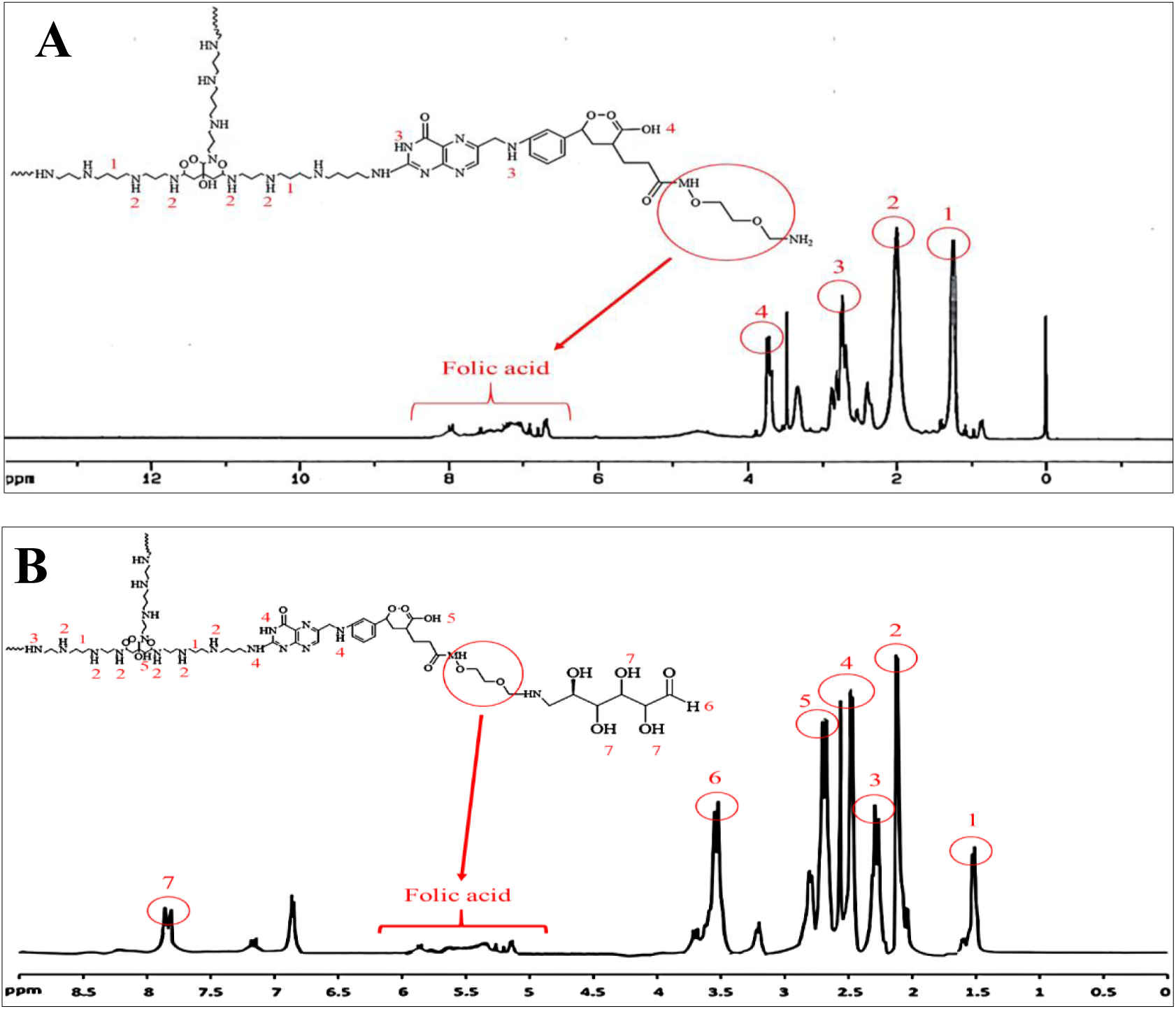
^1^H-NMR spectroscopy **A)** PolySpermine-PEG-FA (PSPF) and **B)** PolySpermine-PEG-FA-Glu (PSPFG).

### 3.2. FT-IR spectroscopy

FT-IR spectroscopy was employed to identify the functional groups of the spermine, polyspermine, and PSPF copolymers. The FT-IR spectrum of spermine exhibited characteristic absorption bands, confirming its polyamine structure (**Figure 5 A**). The broad peak at 3470 cm^-1^ corresponded to the N–H stretching vibrations of the primary (NH_2_) and secondary (NH) amine groups. A sharp band at ∼2850 cm^-1^ was assigned to C–H stretching in methylene (CH_2_) groups. The region between 1600 and 1650 cm^-1^ represented N–H bending in primary amines, whereas the peak at 1530 cm^-1^ represented N–H bending in secondary amines. The absorption near 1150 cm^-1^ was attributed to C–N stretching, and the sharp peak at approximately 1470 cm^-1^ corresponded to CH_2_ bending vibrations.

**Figure 5.**
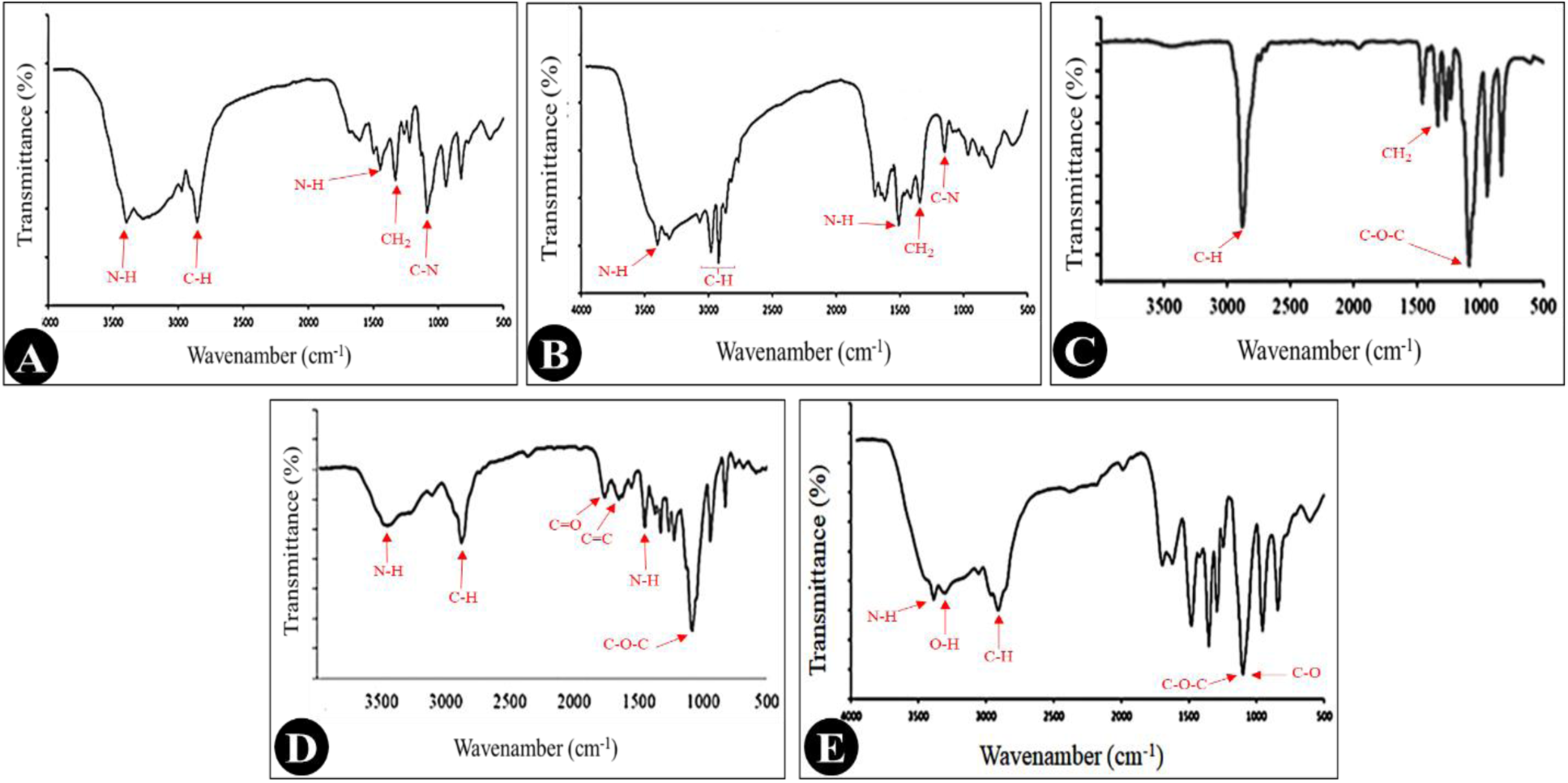
FT-IR spectra of **A)** spermine, B) polyspermine, **C)** PEG, **D)** PSPF, and **E)** PSPFG

Polyspermine displayed spectral features similar to those of spermine, with variations in peak intensity and broadening due to its polymeric nature (**Figure 5 B**). The N–H stretching band of primary and secondary amines appeared at 3400 cm^-1^, whereas the C–H stretching bands of methylene groups were observed between 2850 and 2950 cm^-1^. The band at ∼1480 cm^-1^ corresponded to N–H deformation, and the peaks at 1250 cm^-1^ and 1470 cm^-1^ confirmed C–N stretching and CH_2_ bending, respectively.

The FT-IR spectrum of PEG (**Figure 5 C**) showed a strong absorption band between 2880 and 2900 cm^-1^, corresponding to symmetric and asymmetric CH_2_ stretching vibrations, confirming its polyethylene backbone. A bending band for CH_2_ groups appeared between 1460 and 1480 cm^-1^. The most prominent peak, located between 1100 and 1150 cm^-1^, was attributed to the C–O–C stretching vibrations of the ether linkages.

For the polyspermine-PEG-FA copolymer (**Figure 5 D**), the broad band at 3300--3500 cm^-1^ indicated N–H stretching from polyspermine amines. Methylene C–H vibrations (∼2900 cm^-1^) were present in all three components. The sharp peak at 1750 cm^-1^ corresponded to the carbonyl (C=O) group of folic acid, whereas aromatic C=C stretching was observed near 1650 cm^-1^. The strong band at 1100–1200 cm^-1^ was assigned to C–O–C ether vibrations from PEG. The bands between 1450 and 1500 cm^-1^ were associated with N–H conformational changes and aromatic nitrogen-containing rings of folic acid.

Following glucose conjugation (**Figure 5 E**), a broad peak at ∼3370 cm^-1^ indicated an increase in the hydroxyl group content. The increased intensity of the 2900 cm^-1^ band was consistent with additional methylene groups from glucose. A new band at 1100–1120 cm^-1^, corresponding to C– O stretching in glucose, overlapped with the PEG ether region. Spectral modifications between 1500 and 1650 cm^-1^ were attributed to hydrogen bonding interactions between glucose hydroxyl groups and polyspermine amines. These spectral changes collectively confirmed the successful grafting of glucose onto the nanocapsule surface.

### 3.3. TGA and DTG curves

TGA and DTG were employed to investigate the thermal stability of polyspermine and its modified derivatives (**Figure 6**). The TGA profile of polyspermine (**Figure 6 A**) revealed the onset of degradation at ∼200-300°C, indicative of its high thermal sensitivity in the pure state. The mass loss progressed continuously, with almost complete decomposition occurring at ∼600°C, leaving only trace amounts of thermally stable carbonaceous residues.

**Figure 6.**
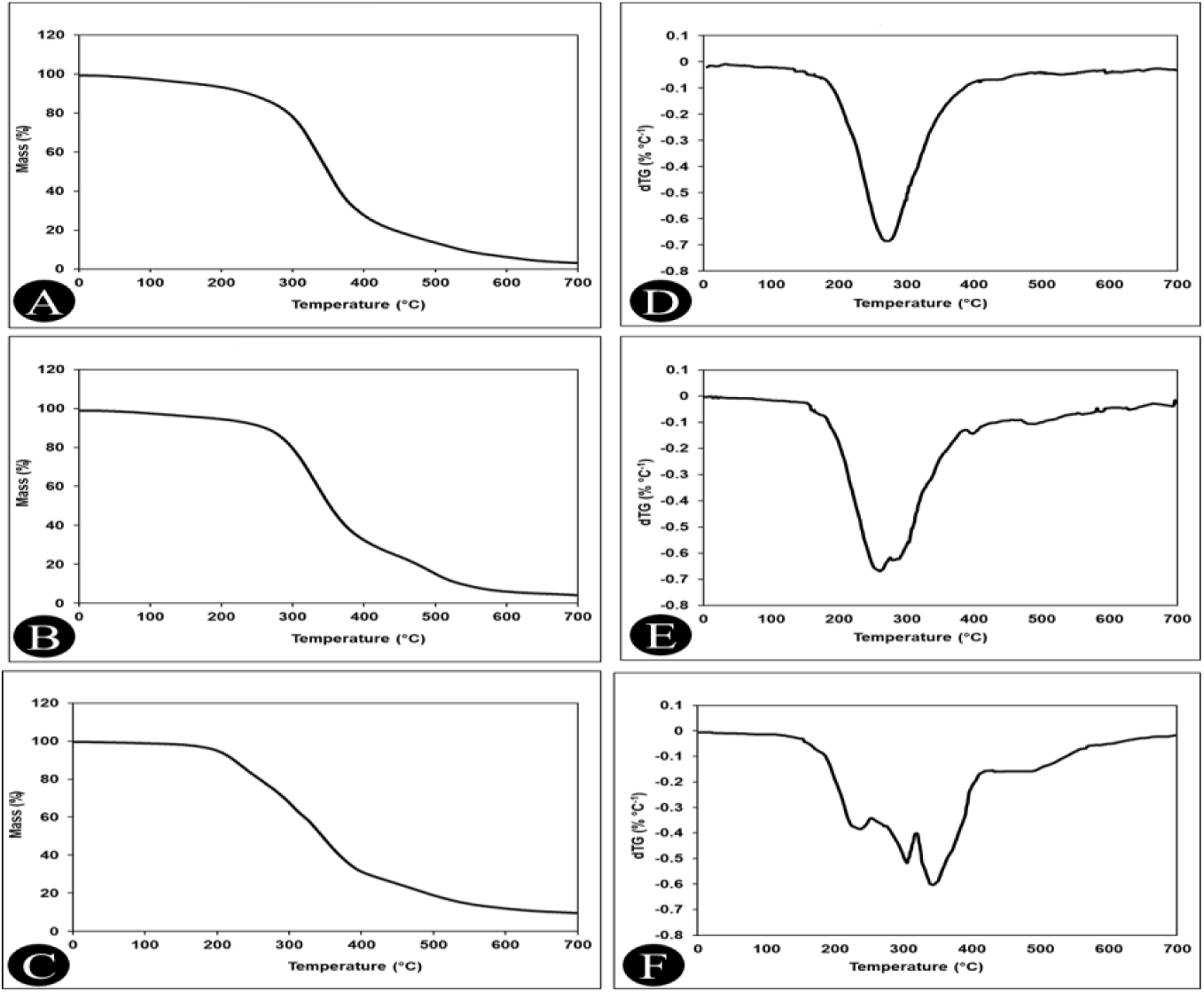
TGA (**A**, **B**, and **C**) and DTG (**D**, **E**, and **F**) curves for PolySpermine, PSPF, and PSPFG, respectively.

In contrast, polyspermine-PEG-FA (PSPF) (**Figure 6 B**) exhibited markedly enhanced thermal stability, initiating degradation at higher temperatures. The weight loss occurred in two distinct stages: (i) 200-350°C, attributed to the decomposition of amino moieties, and (ii) 350-500°C, associated with PEG-FA segment degradation. This multistep behavior reflects the increased molecular complexity of the modified polymer. The PSPF-glucose (PSPFG) sample (**Figure 6 C**) presented the most complex TGA profile, with three separate degradation stages: initial low-temperature weight loss, similar to previous samples, followed by PEG-FA group decomposition, and finally glucose degradation at ∼400-500°C. PSPFG exhibited the highest overall thermal stability among the three materials.

DTG analysis further elucidated the degradation kinetics. Polyspermine (**Figure 6 D**) showed a sharp, intense peak at 250-300°C, indicating rapid mass loss within this range. In PSPF (**Figure 6 E**), broader and less intense peaks were observed across multiple temperature ranges, suggesting stepwise decomposition of different structural segments and a greater diversity of functional groups. PSPFG (**Figure 6 F**) displayed an additional distinct peak at ∼450°C, confirming glucose decomposition and validating the successful chemical modification. A quantitative comparison of the TGA and DTG results revealed that the thermal stability increased with each modification step. This improvement is likely due to stronger covalent linkages, increased molecular weights, and the formation of more intricate polymer architectures, which collectively retard thermal degradation.

### 3.4. Morphological characterization of PSPFG100/DNA nanocapsules

**Figure 7 A** and **B** present TEM and SEM images of PSPFG100/DNA nanocapsules, respectively, clearly illustrating their structure and morphology. In the TEM image, the nanocapsules appeared as spherical to slightly elliptical particles, predominantly within the 200-300 nm range, with a relatively narrow size distribution. The high contrast in the TEM micrograph suggests a suitable electron density, likely resulting from the presence of DNA and the PSPFG polymer. SEM images further confirmed that the nanocapsules possessed a smooth and uniform surface, a desirable feature for drug delivery applications. The particle dimensions fall within the optimal range for traversing biological barriers and reaching target cells. The consistent particle size and morphology reflect the controlled and optimized synthesis process, whereas the minimal aggregation observed in both images indicates good colloidal stability. Collectively, these morphological and structural characteristics position PSPFG100/DNA nanocapsules as promising platforms for the controlled release of therapeutic agents.

**Figure 7.**
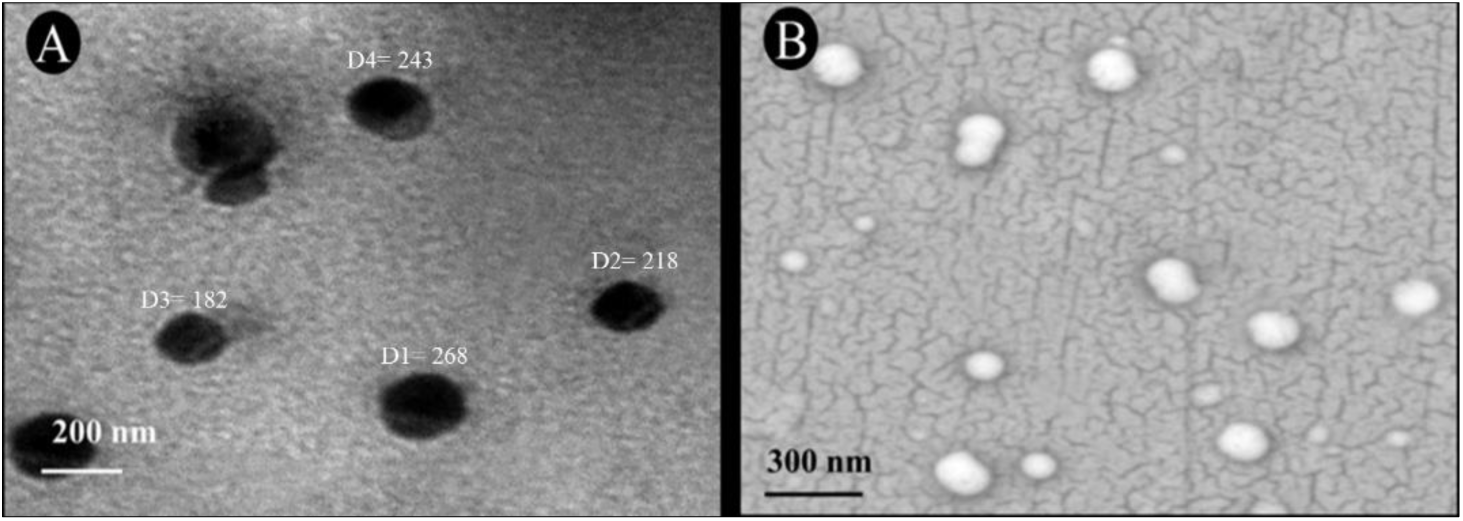
**A)** TEM and **B)** SEM images of PSPFG100/DNA nanocapsules.

### 3.5. Zeta potential and size of the nanocapsules

The results of DLS analysis and zeta potential (Figure 8 and **Table 2**) show that the size of the PSPF nanocapsules is approximately 163 nm, with a standard deviation of ±7.2, and these particles have a negative zeta potential of −6.2, with a standard deviation of ±0.5, which indicates good colloidal stability (**Figure 8 A** and **F**). When glucose was replaced with folic acid, the size of the PSPG nanocapsules changed to 182 nm (±10.2), and the positive zeta potential was +2.7 (±0.7) (**Figure 8 B** and **G**). The different results of these two types of nanocapsules are most likely due to differences in their chemical composition. This confirmed the difference in the nanocapsules. On the other hand, nanocapsules containing both glucose and folic acid (PSPFG) with a size of 211 nm (±15) and a zeta potential of −2.2 (±0.6) showed intermediate properties (**Figure 8 C** and **H**), which indicated the presence of both chemical compounds in the resulting nanocapsules. After DNA loading, the size of the PSPF100/DNA nanocapsules increased to 208 nm (±12), and their zeta potential reached −7.3 (±0.4) (**Figure 8 D** and **I**), indicating the effect of DNA loading on the size and surface charge of the nanocapsules. PSPFG100/DNA nanocapsules also presented the largest size and greatest change in surface charge, with a size of 265 nm (±18) and a zeta potential of +1.7 (±0.7) (**Figure 8 E** and **J**).

**Figure 8.**
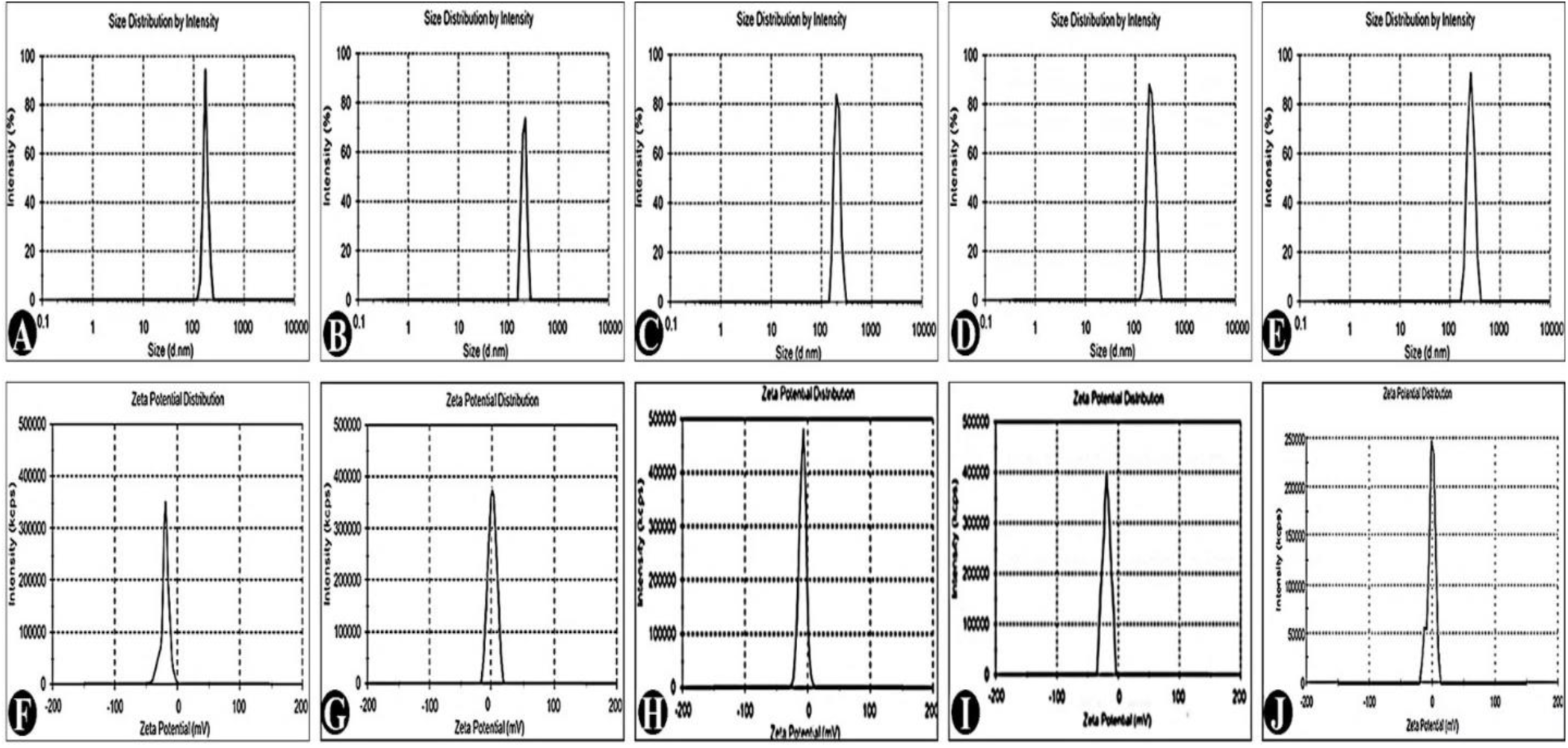
**A, B, C, D, and E)** Size of PSPF, PSPG, and PSPGF nanocapsules and PSPF100/DNA, and PSPFG100/DNA nanocapsules (respectively). **F, G, H, I, and J)** Zeta potentials of PSPF, PSPG, and PSPGF nanocapsules and PSPF100/DNA and PSPFG100/DNA nanocapsules, respectively.

**Table 2.** Size and zeta potential of PSPF, PSPG, and PSPGF nanocapsules and PSPF100/DNA and PSPFG100/DNA nanocapsules.

|  | Size<br>(nm) | Standard<br>deviation | Zeta potential<br>(mV) | Standard<br>deviation | PDI | Standard<br>deviation |
| --- | --- | --- | --- | --- | --- | --- |
| PSPF nanocapsules | 163 | $\pm 7.2$ | - 6.2 | $\pm 0.5$ | 0.23 | $\pm 0.01$ |
| PSPG nanocapsules | 182 | $\pm 10.2$ | + 2.7 | $\pm 0.7$ | 0.27 | $\pm 0.02$ |
| PSPFG nanocapsules | 211 | $\pm 15$ | - 2.2 | $\pm 0.6$ | 0.34 | $\pm 0.06$ |
| PSPF100/DNA nanocapsules | 208 | $\pm 12$ | - 7.3 | $\pm 0.4$ | 0.52 | $\pm 0.05$ |
| PSPFG100/DNA nanocapsules | 265 | $\pm 18$ | + 1.7 | $\pm 0.7$ | 0.66 | $\pm 0.08$ |

PDI is a critical parameter that describes the molecular mass distribution and size uniformity of nanocapsules [36]. Lower PDI values indicate a more consistent particle size distribution, whereas higher values suggest polydisperse samples. The PDI results for the nanocapsules examined in this work are as follows:

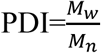

(M_w_= weight average molecular weight, M_n_= number average molecular weight)

The following formulas were used to calculate the M_W_ and M_n_ values:

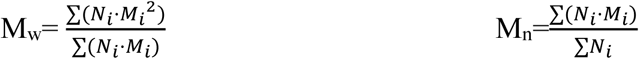

where N_i_ is the number of particles and M_i_ represents the mass of the particles.

The mean diameter and PDI are essential parameters because they determine the transport of nanocapsules *in vivo*. The PDI value can vary from 0.01 (monodisperse particles) to 0.5–0.7, whereas a broad particle size distribution of formulations has been shown with a PDI value > 0.7 [37]. In this study, the PDIs for PSPF, PSPG, and PSPFG nanocapsules were 0.23 (±0.01), 0.27 (±0.02), and 0.34 (±0.06), respectively, indicating their relatively uniform size distributions, but after DNA loading, the PDI values for PSPF100/DNA and PSPFG100/DNA nanocapsules increased to 0.52 (±0.05) and 0.66 (±0.08), respectively, indicating a broadening of the particle size distribution after DNA loading. These changes could be due to the complex interactions between the nanocapsules and the DNA molecules (**Table 2**).

### 3.6. DNA release patterns from PSPG, PSPF, and PSPFG nanocapsules

The DNA loading efficiencies of the PSPG, PSPF, and PSPFG nanocapsules were 43.6%, 40.45%, and 42.33%, respectively. DNA release studies under neutral (pH = 7.4) and acidic (pH = 5.5) conditions, compared with the PEI100/DNA complex, revealed that at pH = 7.4, the PEI100/DNA complex exhibited markedly faster release, particularly in the presence of high dextran concentrations. At 100 μg/mL dextran, the DNA release rates were 64.96% for PEI100/DNA, 43.55% for PSPG, 59.33% for PSPF, and 50.11% for PSPFG (**Figure 9 A**). At pH = 5.5, the release rates of all the nanocapsules and the PEI100/DNA complex increased significantly compared with those under neutral conditions, and no significant differences were observed among the four systems (**Figure 9 B**). The incorporation of folic acid in PSPF and PSPFG nanocapsules enhanced DNA release at both pH values, likely due to electrostatic repulsion between the negatively charged folic acid and the DNA backbone. Overall, the pH-responsive behavior of PSPG, PSPF, and PSPFG nanocapsules, characterized by accelerated release in acidic media, suggests the potential for improved gene or drug delivery to tumor tissues, which typically exhibit a slightly acidic microenvironment.

**Figure 9.**
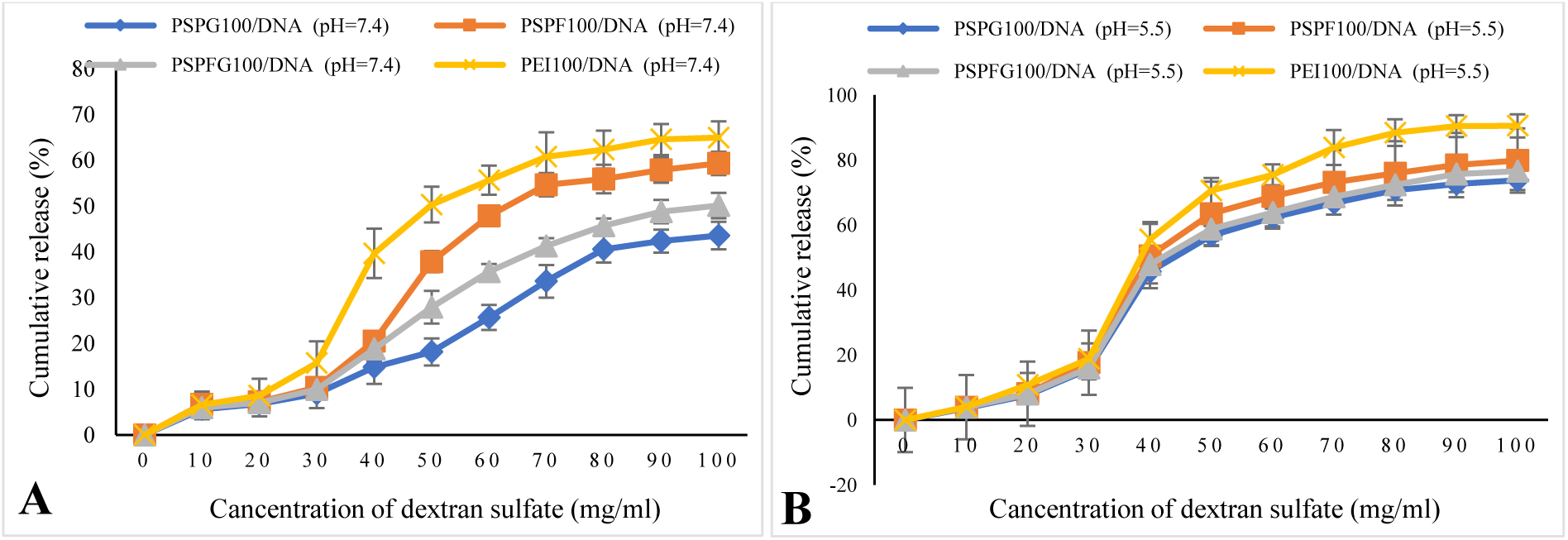
DNA release patterns from PSPG, PSPF, PSPFG, and PEI nanocapsules at different pH values: **A)** pH=7.4 and **B)** pH=5.5.

### 3.7. Biocompatibility assay of PSPG, PSPF, and PSPFG nanocapsules

#### 3.7.1. MTT Assay

One of the most crucial requirements for the use of nanocapsules in clinical settings is their biocompatibility [38]. Thus, assessing the cytotoxicity of nanocarriers is necessary. Even at higher concentrations (100 μg/ml), the viability of AGS cells was above 80%, indicating that PSPG, PSPF, and PSPFG nanocapsules were highly biocompatible with AGS cancer cells, according to the MTT test results (**Figure 10 A, B** and **C**). DNA encapsulation within PSPFG nanocapsules led to a reduction in the percentage of viable cells; however, this decrease was not statistically significant, indicating the high biocompatibility of these nanocapsules and their minimal adverse effects on cell survival, which supports their suitability as carriers for drug or gene delivery (**Figure 10 D** and **Figure 11**). PEI and PEI100/DNA copolymers exhibited significant cytotoxicity toward AGS cells at concentrations exceeding 10 μg/mL (**Figure 10 E** and **F**). The data from this study demonstrated that increasing the PEI100/DNA concentration from 5 μg/mL to 100 μg/mL resulted in a decrease in AGS cell viability from 84.45% to 11.34% (**Figure 11**).

**Figure 10.**
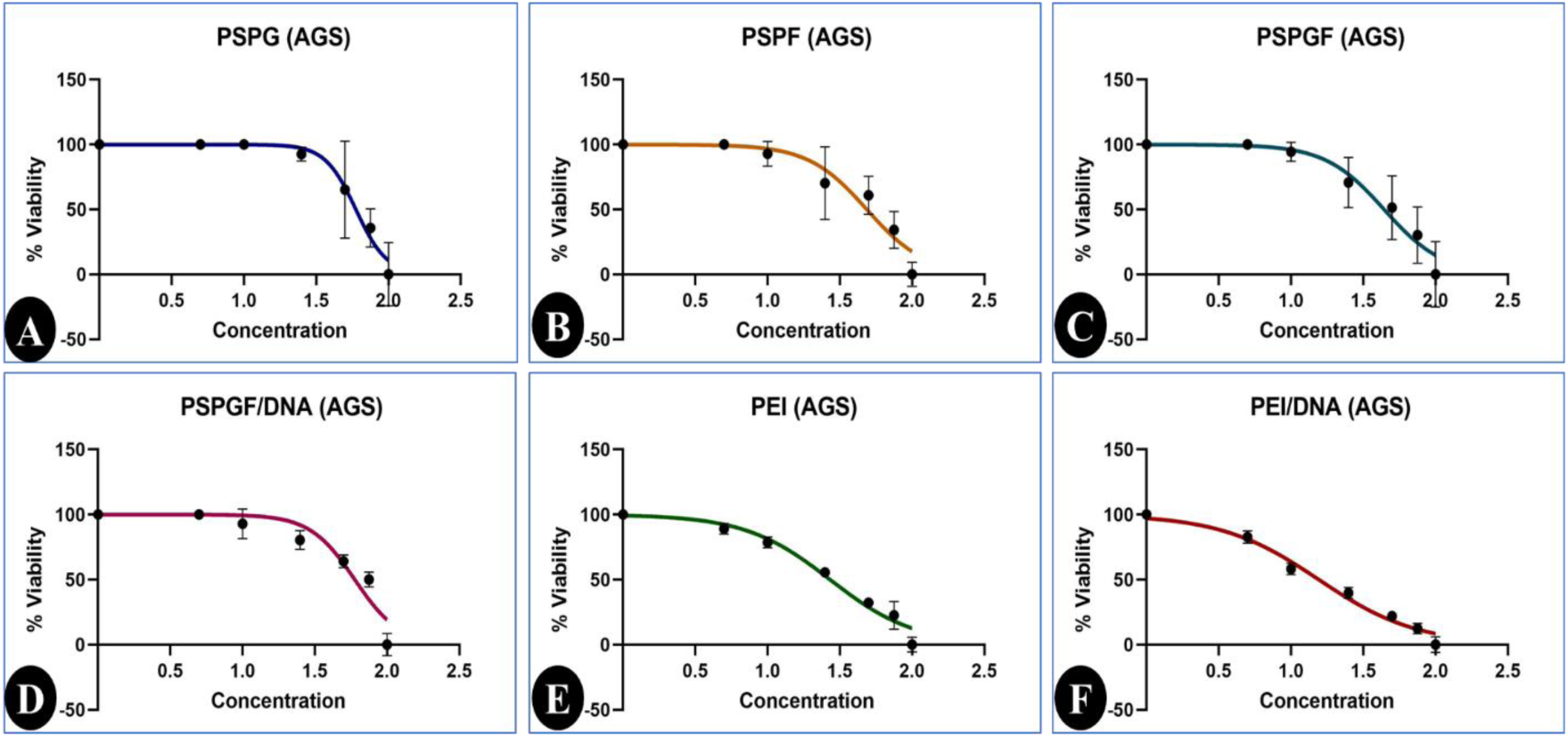
Comparison of the average effects of **A)** PSPG, **B)** PSPF, **C)** PSPFG, **D)** PSPFG100/DNA, **E)** PEI, and **F)** PEI100/DNA copolymers on the viability of AGS cell lines.

**Figure 11.**
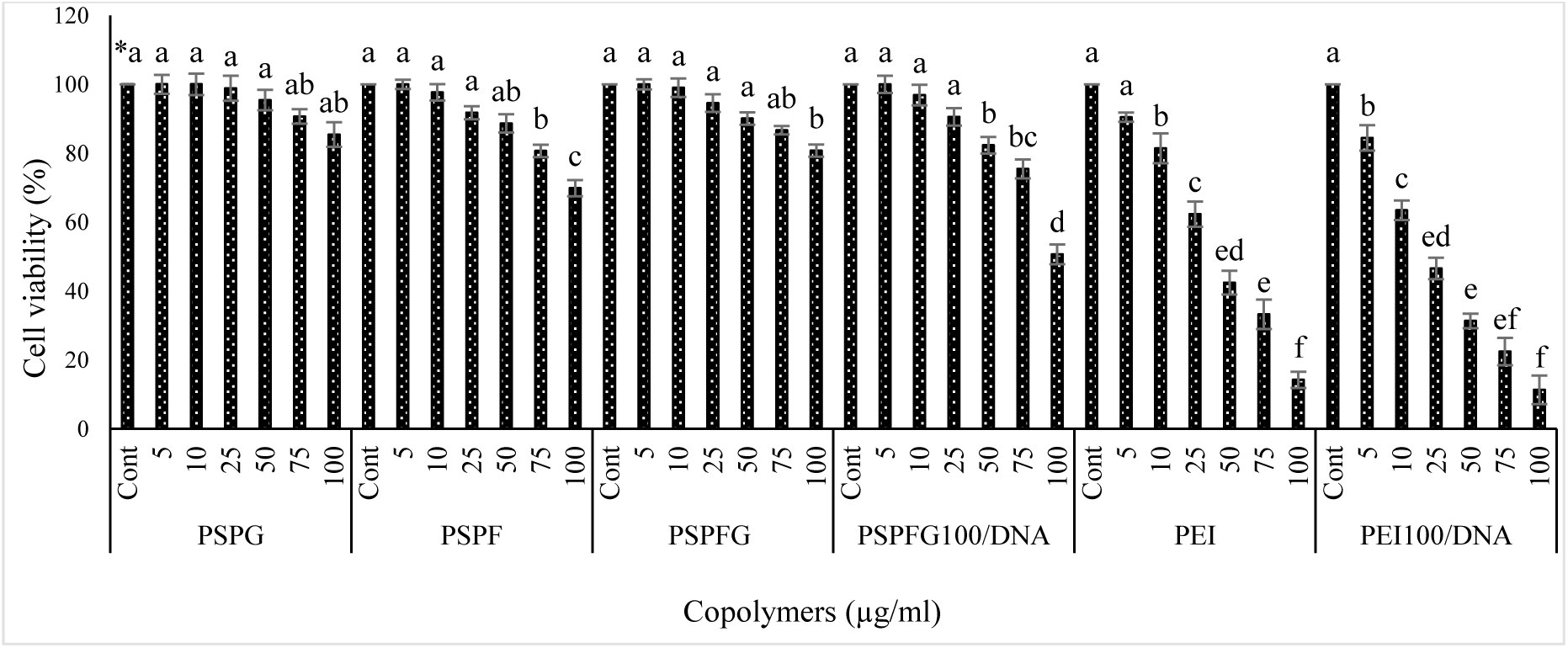
Toxicity of PSPF, PSPG, PSPGF, PEI, PSPFG100/DNA, and PEI100/DNA nanocapsules (at different concentrations) to AGS cells. * Lowercase letters (a, b, c) above the bars or data points represent statistical groupings based on ANOVA followed by Duncan’s multiple range test. Identical letters denote no statistically significant difference, whereas differing letters reflect a significant difference at p < 0.05.

#### 3.7.2. Apoptosis

Targeted drug delivery significantly enhances apoptosis in cancer cells, making the use of targeted nanocapsules highly advantageous [39]. After the IC_50_ values of the tested compounds were determined, their capacities to induce apoptosis and necrosis were evaluated. The treatment with PEI100/DNA complexes resulted in the greatest necrotic cell population (33.56%), which was notably greater than that observed with the other formulations. This suggests that while PEI100/DNA is effective at inducing cell death, its mechanism may favor necrosis over apoptosis compared with targeted nanocapsules, highlighting differences in cytotoxic pathways among the tested nanocarriers (**Figure 12** and **Figure 13 C**). Furthermore, previous studies reported that the elevated percentage of necrotic cells observed in the control group may be attributed to the rapid proliferation of the AGS cell line and subsequent nutrient competition (**Figure 12** and **Figure 13 A**).

**Figure 12.**
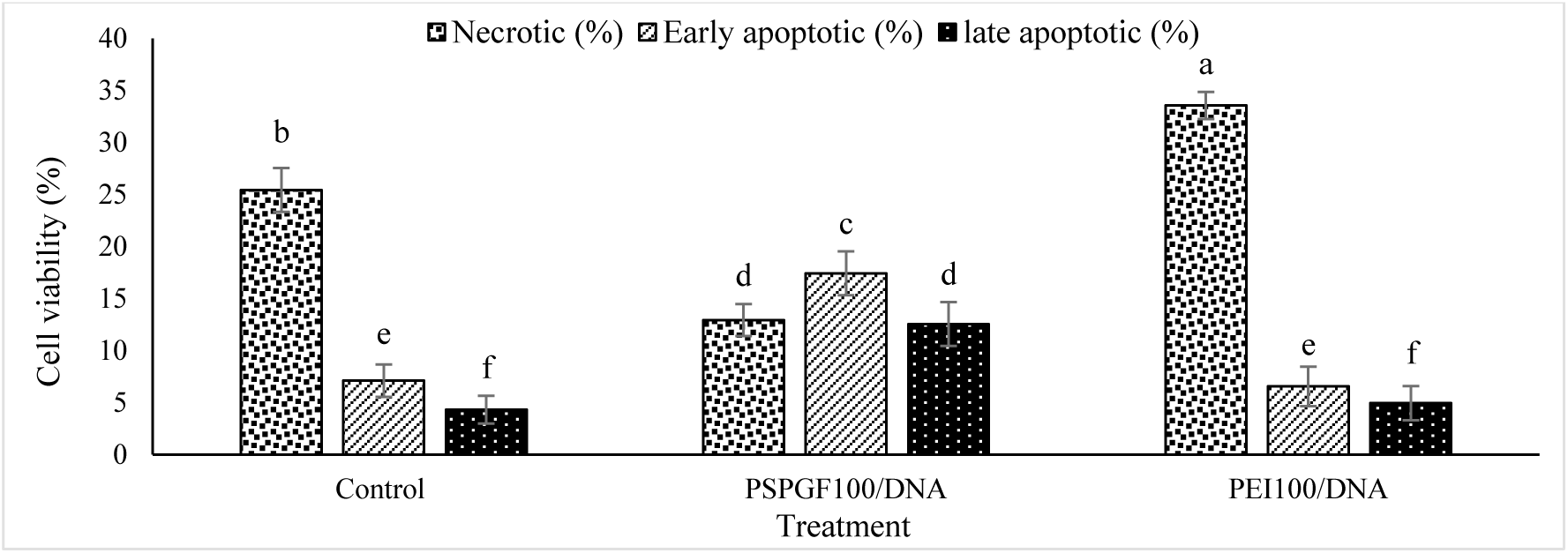
Percentage of AGS cells located in the necrotic, pre and post-apoptosis stages affected by the calculated IC_50_ concentrations of PSPFG100/DNA, PEI100/DNA nanocapsules and the control treatment. * Lowercase letters (a, b, c) above the bars or data points represent statistical groupings based on ANOVA followed by Duncan’s multiple range test. Identical letters denote no statistically significant difference, whereas differing letters reflect a significant difference at p < 0.05.

**Figure 13.**
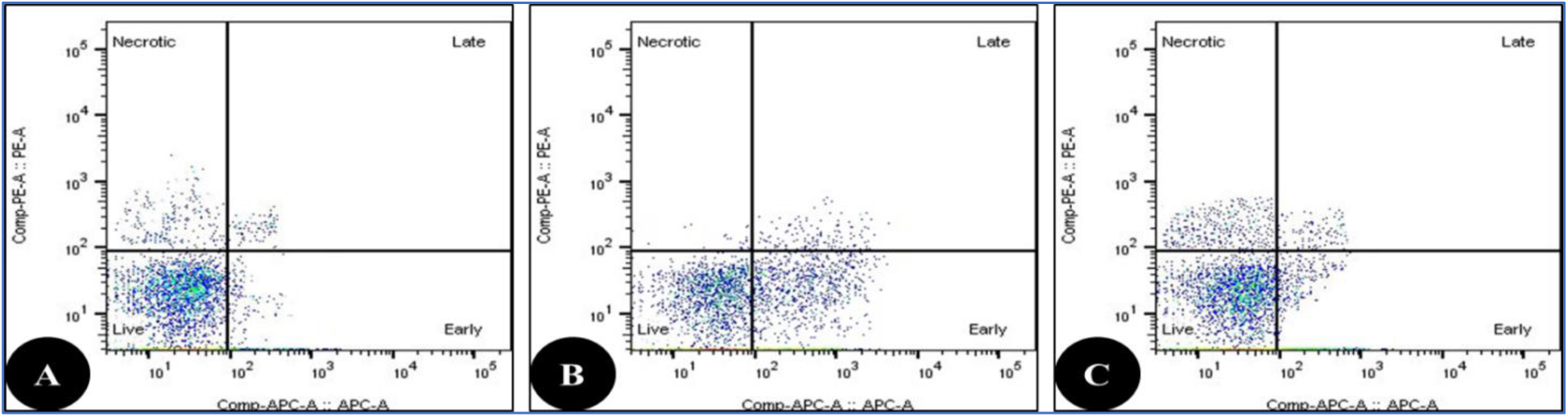
Schematic flow cytometry image of AGS cells in the necrotic, pre- and - phases under the influence of the calculated IC_50_ concentrations of **A)** the control treatment, **B)** PSPFG100/DNA and **C)** PEI100/DNA.

The low percentage of necrotic cells observed following treatment with PSPFG100/DNA nanocapsules suggests minimal cytotoxic side effects from the nanocapsules, alongside a significant induction of apoptosis. Notably, the PSPFG100/DNA group presented the greatest proportion of cells in the early and late stages of apoptosis, at 17.43% and 12.56%, respectively (**Figure 12** and **Figure 13 B**). These findings demonstrate the effective role of nanocapsule carriers in promoting programmed cell death, highlighting their potential as safe and efficient delivery systems.

### 3.8. DNA protection assay of PSPFG/DNA nanocapsules

Agarose gel electrophoresis was utilized to assess the capacity of PSPFG/DNA nanocapsules to neutralize the negative charge of DNA and protect it from nuclease-mediated degradation. The results confirmed that PSPFG nanocapsules effectively neutralized the negative charge of DNA and provided significant protection against plasma restriction enzymes (**Figures 14 and 15**). Owing to its intrinsic negative charge, free DNA migrates toward the positive electrode during electrophoresis; however, when complexed with cationic polymers, DNA mobility is hindered as a result of charge neutralization. As shown in **Figure 14**, PSPFG nanocapsules exhibited a strong ability to neutralize the negative charge of DNA, thereby reducing electrophoretic migration.

**Figure 14.**
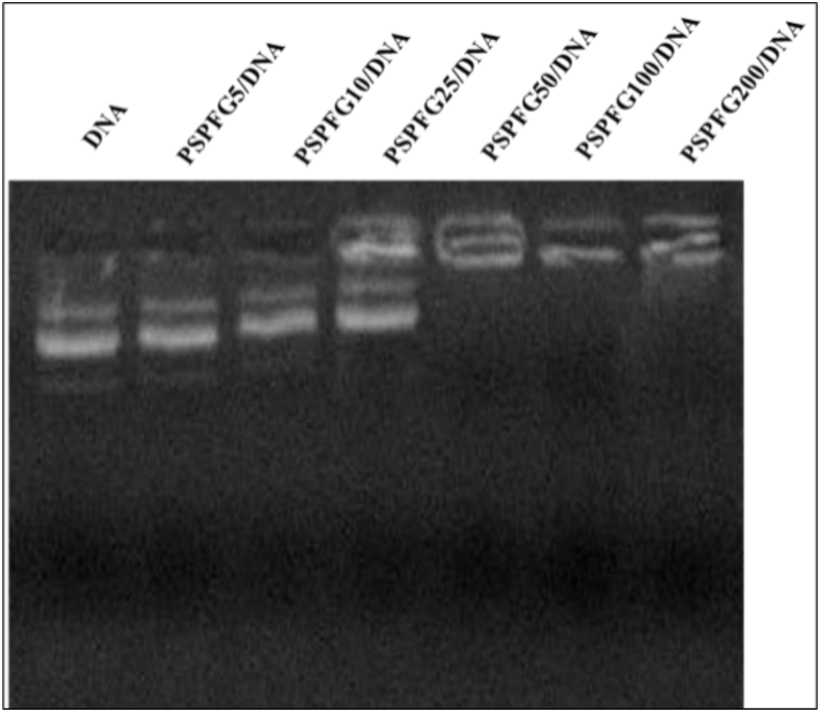
Effect of PSPFG nanocapsules on the neutralization of the negative charge of DNA.

**Figure 15.**
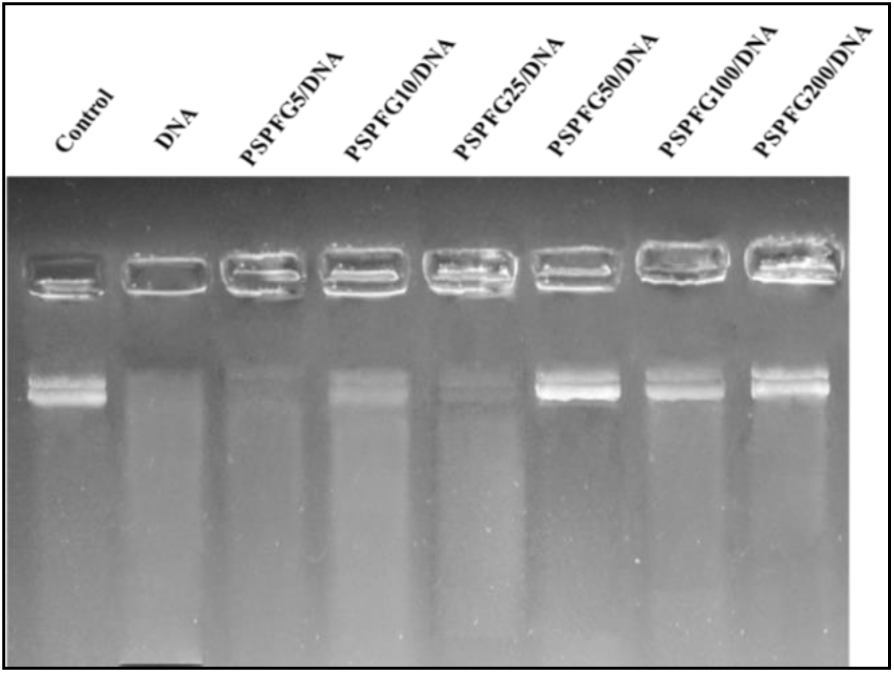
Protective effect of PSPFG nanocapsules on DNA against plasma degradation.

A concentration of 50 μg of PSPFG/DNA nanocapsules was required to completely neutralize 5 μg of DNA. As shown in **Figure 15**, DNA encapsulated within PSPFG/DNA nanocapsules remained intact following incubation with human blood plasma, whereas free (naked) DNA exhibited no visible band on the agarose gel after plasma treatment, indicating fragmentation into smaller pieces.

### 3.9. DNA transfer efficiency of PSPFG nanocapsules to AGS cells

The gene delivery efficiency of PSPFG nanocapsules encapsulating the pEGFP-N1 plasmid into AGS cells was evaluated via flow cytometry and fluorescence microscopy. The results demonstrated effective DNA release and cellular uptake mediated by these nanocapsules (**Figure 16**). Notably, increasing the PSPFG polymer concentration within the nanocapsules significantly improved the gene transfection efficiency; PSPFG50/DNA formulations achieved a gene expression level of 53.37%, substantially exceeding the 18.36% observed with PSPFG5/DNA nanocapsules. No statistically significant differences were detected among the PSPFG50/DNA, PSPFG100/DNA, and PSPFG200/DNA groups (**Figure 17**), indicating a saturation plateau in gene transfer efficiency at higher polymer concentrations.

**Figure 16.**
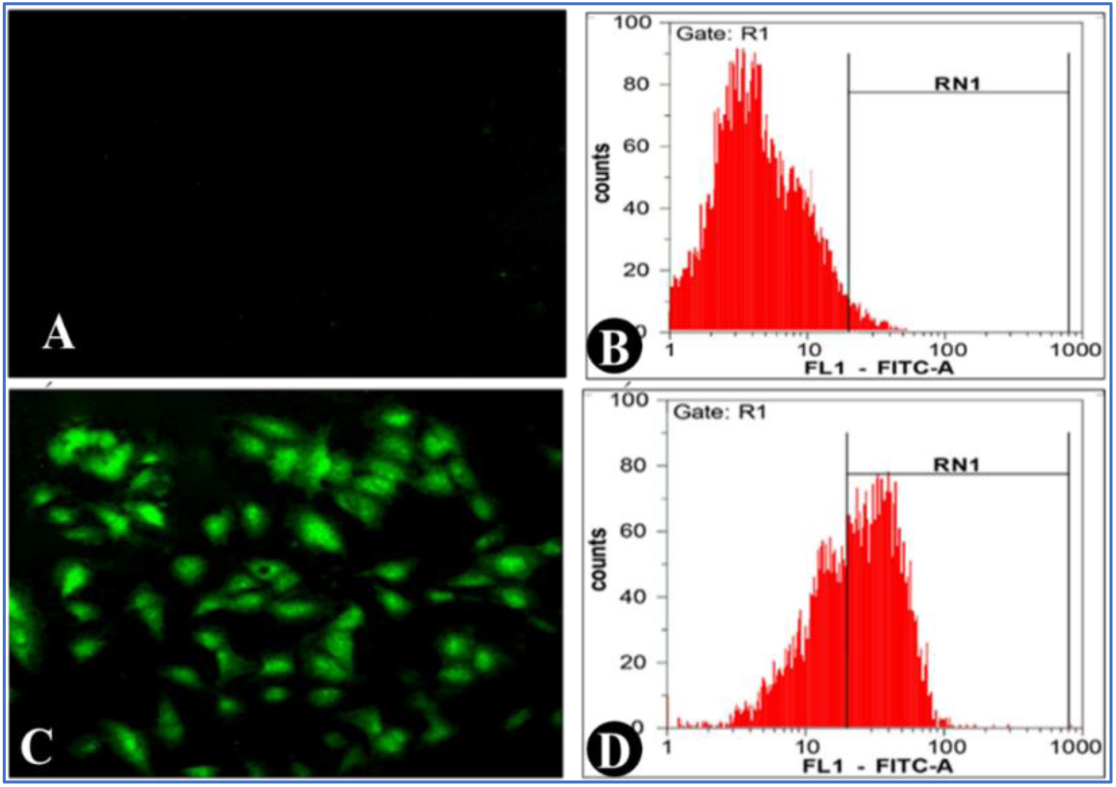
Flow cytometry (**B** and **D**) and fluorescence microscopy (**A** and **C**) images of control cells (**A** and **B**) and cells treated with PSPFG100/DNA nanocapsules (**C** and **D**).

**Figure 17.**
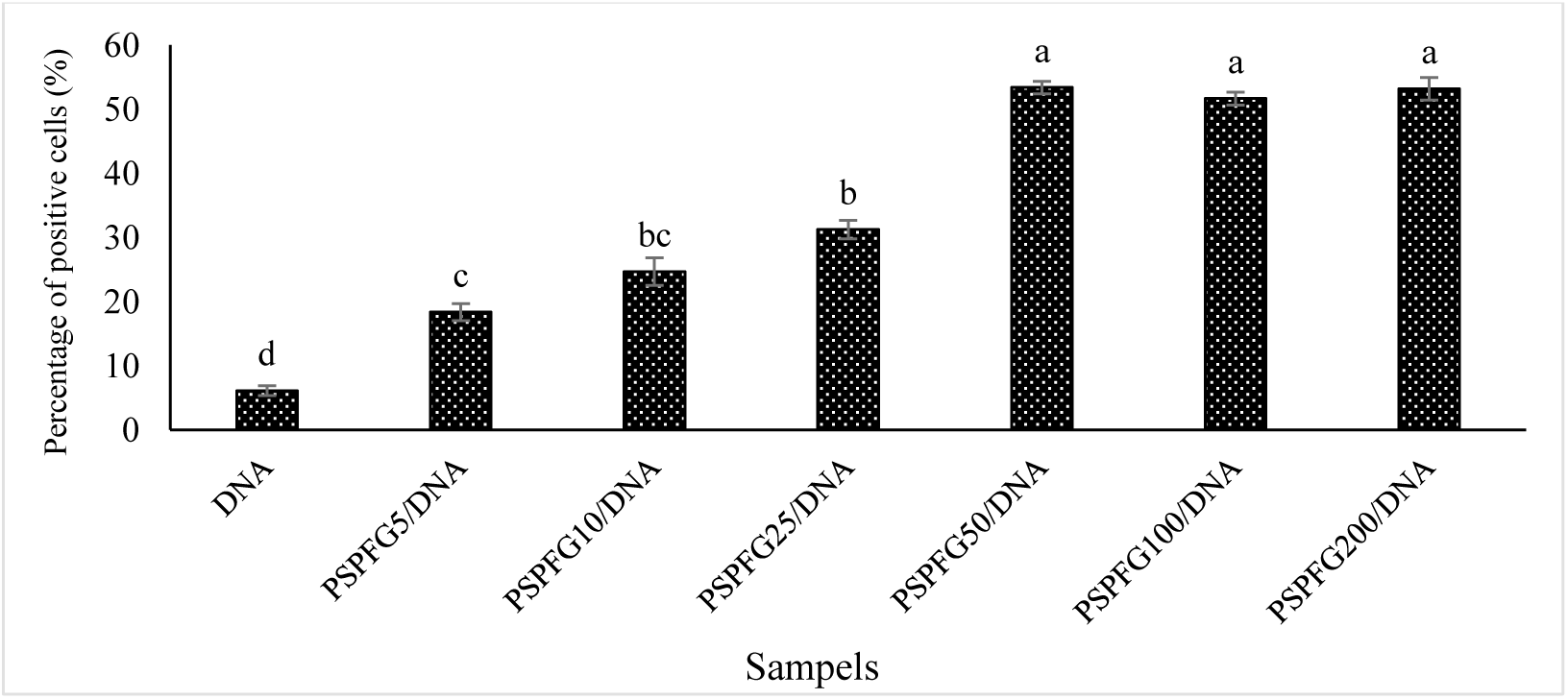
Comparison of the average gene transfer efficiency of PSPFG/DNA nanocapsules in AGS cells. * Lowercase letters (a, b, c) above the bars or data points represent statistical groupings based on ANOVA followed by Duncan’s multiple range test. Identical letters denote no statistically significant difference, whereas differing letters reflect a significant difference at p < 0.05.

## 4. Discussion

Copolymer-based nanoparticles have emerged as promising gene delivery vectors because of their biocompatibility, tunable surface properties, and ability to encapsulate genetic materials [40]. Cationic polymers, in particular, enable electrostatic interactions with negatively charged DNA, facilitating cellular uptake through endocytosis [41]. The gene transfer process using cationic polymers can help reduce the side effects associated with common treatments, such as chemotherapy, and thus increase the hope of improving the quality of life of cancer patients [42].

PEI has been widely studied for its high transfection efficiency, although its toxicity limits its clinical application [43]. Modifications such as ligand conjugation and N/P ratio optimization have been explored to increase specificity and reduce adverse effects [23, 44]. Because of its toxicity, the PEI polymer has not been widely used in clinical settings, despite its high gene transfer efficiency. Numerous investigations into the mechanism of the toxicity of cationic polymers, including PEI, to cancer cells have been carried out [45]. Natural polyamines such as polyspermine offer a biocompatible alternative, with intrinsic DNA-binding capacity and favorable structural characteristics [46]. Targeted delivery using ligands such as folic acid or glucose further improves specificity toward cancer cells [12]. On the basis of the abovementioned cases and the unique and widely used properties of folic acid and glucose in identifying cell surface receptors and targeted drug delivery [47], in this study, folic acid and glucose were used as cell surface receptors for targeted DNA transfer to AGS cancer cells by synthesizing copolymer nanoparticles using polyspermine and PEG.

According to the results obtained from ^1^HNMR, FTIR, TGA and DTG, the structure and nature of various polymers, such as polyspermine, PEG with glucose or folic acid, and the correct synthesis of the polyspermine-PEG/folic acid copolymer were confirmed **(Figures 4, 5 and 6)**. In 2022, Wadhawan et al. synthesized PLA-PEG copolymers conjugated with folic acid and confirmed their functionalization via ^1^H-NMR, identifying aromatic folate proton signals at ∼6.6 and 7.5 ppm, which is consistent with folic acid signatures in our PSPF copolymer at 6–8 ppm. Additionally, the emergence of new ^1^H-NMR peaks upon glucose addition parallels other multifunctional copolymer systems where sugar moieties introduce characteristic resonances in the same region [48]. In a 2025 study, Mohajeri et al. synthesized polyspermine-PEG-glucose nanocapsules for targeted gene delivery and used FT-IR spectroscopy to confirm their functional groups. Their spectra showed broad N–H stretching peaks at 3300–3500 cm^-1^ and increased hydroxyl peaks (∼3370 cm⁻¹) after glucose conjugation, closely matching the spectral shifts observed in our PSPF-glucose nanocapsules [28]. On the other hand, Ramalho et al. (2024) developed folic acid-conjugated poly(lactic-co-glycolic acid) (PLGA) nanocapsules loaded with gallic acid for targeted glioblastoma therapy. Their FT-IR spectra confirmed the successful incorporation of folic acid, with distinct peaks at 1750 cm^-1^ (C=O stretching) and 1650 cm^-1^ (aromatic C=C bending), which is consistent with our observations in the FT-IR spectra of PSPF copolymers [49].

The shape and morphology of nanocapsules, as key factors in the targeted gene transfer process, are highly important and significantly affect the efficiency of these delivery systems [50]. Nanocapsules of different shapes and sizes can exhibit their own physical and chemical properties, which directly affect their interactions with biological tissues and cells. TEM and SEM analyses of PSPFG100/DNA nanocapsules revealed well-defined spherical to elliptical particles with sizes predominantly between 200 and 300 nm, confirming a relatively narrow size distribution (**Figure 7**). The high contrast in the TEM images suggests effective incorporation of both DNA and polymer, whereas the SEM images reveal smooth and uniform nanocapsule surface features favorable for cellular uptake and controlled drug release. The observed uniformity in morphology and limited particle aggregation indicate that an optimized synthesis process yields stable colloidal nanocapsules suitable for crossing biological barriers and targeted delivery applications. A recent study published in 2024 reported the synthesis of DPLA-PEG-folic acid nanocapsules with an average size of approximately 241 nm. SEM images demonstrated that these nanocapsules possess a uniform spherical morphology and smooth surfaces, closely aligning with our observations of PSPFG100/DNA nanocapsules [51]. These morphological characteristics are essential for efficient drug and gene delivery, as they contribute to favorable cellular uptake and stability. These findings support the suitability of our nanocapsules for targeted therapeutic applications.

The size and surface charge of the nanocapsules are critical factors influencing the efficiency of targeted gene delivery. These properties affect the cellular uptake, membrane penetration, and overall stability of the delivery system. Optimizing size and charge is essential both scientifically and clinically to increase the precision and effectiveness of gene therapy while minimizing delivery challenges [52]. DLS measurements revealed a substantial increase in nanocapsule size after DNA incorporation, with PSPF100/DNA and PSPFG100/DNA formulations reaching average diameters of 208 nm and 265 nm, respectively. This increase reflects the successful association of DNA with the nanocapsule surface or internal structures. Concurrently, zeta potential measurements revealed pronounced shifts from −6.2 mV to −7.3 mV for PSPF and from −2.2 mV to +1.7 mV for PSPFG-, highlighting changes in surface charge, likely due to DNA binding and rearrangement of the nanocapsule corona. Additionally, the PDI increased from 0.2–0.3 to 0.5–0.6 postloading, indicating a broader size distribution, possibly arising from variable binding efficiencies and structural heterogeneity in the PSPFG100/DNA complexes. Together, these findings confirm effective DNA encapsulation and underscore its significant impact on the physicochemical behavior of the resulting nanosystems (**Figure 8** and **Table 2**). A recent study by Okami et al. (2024) demonstrated the successful synthesis of plasmid DNA-loaded lipid nanocapsules via a one-step microfluidic method, resulting in particles ranging from 95 to 157 nm with PDI values ranging from approximately 0.2-0.3. DNA loading significantly influences the surface charge of nanocapsules, with zeta potentials shifting between −9 and −19 mV [53]. These findings strongly support our observations, confirming that DNA encapsulation alters both the size and physicochemical properties of nanocarriers in a predictable manner.

Gene encapsulation in nanocapsules is influenced by both physical factors, such as size, shape, and surface characteristics, and chemical factors, including coating composition and stability [54]. Smaller nanocapsules and specific shapes tend to enhance cellular penetration and interaction with cell membranes [12]. Environmental conditions such as pH and temperature also play crucial roles, particularly in modulating gene release from nanocapsules [55]. pH affects the electrical charge and binding stability of gene-nanocapsule complexes, where a neutral pH favors efficient gene delivery, whereas extremely acidic or alkaline conditions hinder this process. Importantly, cancer cells typically exhibit a more acidic microenvironment than healthy cells do (pH=4.7), which impacts their metabolism, invasiveness, and immune response [38]. This pH difference not only influences gene delivery efficiency but also presents opportunities for targeted therapeutic strategies by exploiting tumor acidity [56]. Understanding these intertwined factors is essential for optimizing nanocapsule-based gene therapies. Our release studies indicate that DNA from PEI100/DNA nanocapsules is released more rapidly at neutral pH =7.4 than that from PSPG, PSPF, and PSPFG nanocapsules. Under acidic conditions (pH=5.5), however, all formulations, including PEI100/DNA, similarly accelerated DNA release, with no significant differences (**Figure 9**). Notably, PSPF, and PSPFG exhibited greater DNA liberation than PSPG did, suggesting that folic acid plays a key role in facilitating payload release across both pH environments. This pH-sensitive behavior aligns well with the more acidic microenvironment of tumors and highlights the potential of PSPFG nanocapsules for targeted gene delivery in cancerous tissues. A 2024 study by Sharma et al. evaluated pH-responsive, folate-conjugated chitosan nanoparticles loaded with curcumin and reported significantly faster drug release at acidic pH (5.5) than at neutral pH (7.4) due to protonation of the amine groups that swell the polymer matrix [57].

Biocompatibility is essential for the clinical use of micellar carriers, and our MTT results show that PSPG, PSPF, and PSPFG nanocapsules maintain over 80% AGS cell viability even at 100 μg/mL, indicating minimal toxicity (**Figures 10 and 11**). In contrast, the PEI100/DNA complexes triggered severe cytotoxicity, reducing viability to just 11% at the same concentration and primarily inducing necrosis (∼33%). Importantly, PSPFG100/DNA formulations elicited the highest levels of structured apoptosis (∼17% early + ∼13% late-stage), with minimal necrosis, highlighting their potential as safe, efficient gene delivery vehicles with a clear advantage over conventional PEI carriers (**Figures 12 and 13**). In a study by Soltani et al., folic acid- and quercetin-loaded cellulose nanoparticles were developed and tested on AGS cancer cells, where they maintained over 80% cell viability even at high concentrations while still inducing notable levels of apoptosis. These findings are consistent with our results for PSPFG-based nanocapsules, further supporting the effectiveness of folate-functionalized polymer carriers in achieving targeted therapeutic delivery with minimal cytotoxicity [58].

However, the findings of this study demonstrated that PSPG, PSPF, and PSPFG nanocapsules are significantly less harmful than PEI is (**Figures 10 and 11**). The observed reduction in toxicity may be attributed to PEG shielding the positive charge of spermine, given that the polymer’s cytotoxicity is primarily linked to its strong positive charge and interactions with the cell membrane. Studies investigating PEI polymer toxicity across various animal cell lines have demonstrated that nanocapsule-induced cytotoxicity arises from membrane damage mediated by their pronounced positive charge [29]. Covalent conjugation of PEG to PEI polymers represents a key strategy for reducing polymer-associated toxicity. However, a notable limitation of PEGylation is its potential to hinder intracellular DNA release, thereby increasing the risk of DNA degradation within endosomal compartments [23].

Gene delivery is more complex than conventional chemotherapy because of the large size and vulnerability of DNA, requiring protective nanocapsule systems that increase stability and cellular uptake [59]. Cationic polymers play a crucial role by condensing DNA through electrostatic interactions, shielding it from enzymatic degradation and preventing restriction enzyme access, thereby improving delivery efficiency [60]. The results of this study also revealed that the interaction of PSPGF with DNA protects DNA from enzymatic digestion (**Figures 14 and 15**). Agarose gel electrophoresis demonstrated that PSPFG/DNA nanocapsules effectively neutralized the negative charge of the DNA, completely blocking migration at a 50:5 µg ratio, and robustly protected the DNA from degradation in human plasma, where the naked DNA was fully fragmented (**Figure 14**). This finding indicates that the cationic PSPFG coating forms a compact shell around the DNA, preserving its integrity against nucleases. These features are critical for delivering intact gene material *in vivo*, highlighting the potential of PSPFG nanocapsules as protective and efficient gene delivery carriers.

The agarose gel electrophoresis results demonstrated that PSPFG/DNA nanocapsules effectively neutralized the negative charge of the DNA and provided significant protection against nuclease degradation in the plasma (**Figure 15**). The encapsulated DNA remained intact after incubation with plasma enzymes, whereas free DNA was completely degraded. These findings highlight the strong protective ability of PSPFG nanocapsules, underscoring their potential as safe and efficient gene delivery carriers in biological systems (**Figures 16 and 17**). Chandrasekaran and Halvorsen (2020) presented a detailed protocol for analyzing DNA nanostructure stability against nuclease degradation via agarose gel electrophoresis [61]. Their work emphasized how cationic carriers can shield DNA from enzymatic cleavage, which aligns with the protective effects observed in PSPFG/DNA nanocapsules. Therefore, according to the results of similar studies, compression and reduction of the contact surface of DNA by PSPGF nanocapsules prevent the binding of restriction enzymes.

## Conclusion

This study aimed to propose a targeted gene transfer method for gastric cancer cells (AGS) that utilizes the benefits of cationic and biocompatible polymers. These findings demonstrated that combining polyspermine with PEG enhances nanoparticle biocompatibility and facilitates DNA attachment. Coating nanoparticles with PEG improves their solubility in plasma, decreasing immune detection and clearance. The PEG surface was modified with glucose, which is more abundant on cancer cells than on normal cells. Consequently, targeting nanoparticles with folic acid and glucose enhances their delivery to cancer cells while minimizing their uptake in normal cells. The study concludes that spermine is a promising alternative to traditional cationic polymers such as PEI.

## Author contributions

M.R. and S.B. conducted the experiments, analyzed the data, and wrote the original draft. H.Y. and S.M. conceived and designed the research, administered and supervised the project, and M.N. analyzed the gene data and reviewed and edited the manuscript. All the authors read and approved the manuscript.

## Ethics statement

This article does not contain any studies with human participants or animals performed by any of the authors.

## Data availability

The data that support the findings of this study are available from the corresponding author upon reasonable request.

## Consent for publication

Not applicable

## Participating declaration

Not applicable

## Abbreviation list

MTT: 3-(4,5-Dimethyl-2-thiazolyl)-2,5-diphenyl-2H-tetrazolium bromide
DTG: Differential thermal gravimetric
DMEM: Dulbecco’s modified Eagle’s medium
DLS: Dynamic light scattering
EE: Encapsulation efficiency
EPR: Enhanced permeability and retention
EDC: Ethylene dichloride
EDTA: Ethylenediaminetetraacetic Acid
FBS: Fetal bovine serum
FTIR: Fourier transform infrared spectroscopy
Glu: Glucose
GLUTs: Glucose receptors
^1^HNMR: Hydrogen nuclear magnetic resonance spectroscopy
DMSO: Methyl sulfoxide
NPs: Nanoparticles
NHS: N-hydroxysuccinimide
PDI: Polydispersity index
PEG: Polyethylene glycol
PEI: Polyethyleneimine
PLL: Poly-L-lysine
PS: PolySpermine
PSPG: Poly-Spermine-polyethylene glycol-glucose
PSPF: PolySpermine-polyethylene glycol-Folic acid
PVA: Polyvinyl alcohol
PVP: polyvinyl pyrrolidone
SEM: Scanning electron microscopy
TGA: Thermogravimetric analysis
TEM: Transmission electron microscopy

## References

1. Lordick F, Carneiro F, Cascinu S, Fleitas T, Haustermans K, Piessen G, Vogel A, Smyth E. Gastric cancer: ESMO Clinical Practice Guideline for diagnosis, treatment and follow-up☆. Annals of Oncology. 2022;33(10):1005–20.

2. Alsina M, Arrazubi V, Diez M, Tabernero J. Current developments in gastric cancer: from molecular profiling to treatment strategy. Nature Reviews Gastroenterology & Hepatology. 2023;20(3):155–70.

3. Noruzpuor M, Asghari Zakaria R, Zare N, Ebrahimi HA, Parsa H, Bourang S. Green synthesis of metal nanoparticles using aqueous extract of *Moringa oleifera* L. and investigating their antioxidant and antibacterial properties. Applied Chemistry Today. 2024;19(71):283–302.

4. Asghari Zakaria R, Zare N, Ebrahimi HA, Parsa H, Bourang S. Investigating the Anticancer Properties of the Essential Oil and Aqueous Extract of *Moringa oleifera* and its Biosynthesized Metal Nanoparticles on MCF-7 and BT-549 Cell Lines. Iranian Journal of Breast Diseases. 2024;17(1):59–83.

5. Jahazi S, Akbari H. Preparation and characterization of doxorubicin loaded Fe3O4-PEG nanoparticles on AGS and MCF-7 cancer cells. Modares Journal of Biotechnology. 2020;11(2):167–75.

6. Bourang S, Noruzpour M, Jahanbakhsh Godekahriz S, Ebrahimi HAC, Amani A, Asghari Zakaria R, Yaghoubi H. Application of nanoparticles in breast cancer treatment: a systematic review. Naunyn-Schmiedeberg’s Archives of Pharmacology. 2024:1–47.

7. Bahrami B, Hojjat-Farsangi M, Mohammadi H, Anvari E, Ghalamfarsa G, Yousefi M, Jadidi-Niaragh F. Nanoparticles and targeted drug delivery in cancer therapy. Immunology letters. 2017;190:64–83.

8. Bourang S, Jahanbakhsh-Godekahriz S, Asghari-Zakaria R, Parsa-Khankandi H, Noruzpour M. Green synthesis of iron oxide, copper, zinc oxide and silver nanoparticles from aqueous extract of *F. vulgare* and evaluation of their structural and antimicrobial properties. Agricultural Biotechnology Journal. 2024;16(3):61–88.

9. Cring MR, Sheffield VC. Gene therapy and gene correction: targets, progress, and challenges for treating human diseases. Gene therapy. 2022;29(1):3–12.

10. Mohajeri S, Dashti S, Noruzpour M, Bourang S, Yaghoubi H. Design and preparation of PLA-chitosan-PEG-glucose copolymer for combined delivery of Paclitaxel and siRNA. Discover Applied Sciences. 2025;7(8):801.

11. Ahmadi-Nouraldinvand F, Bourang S, Azizi S, Noori M, Noruzpour M, Yaghoubi H. Preparation and characterization of multi-target nanoparticles for co-drug delivery. Medicine in Drug Discovery. 2024;21:100177.

12. Kashani GK, Naghib SM, Soleymani S, Mozafari M. A review of DNA nanoparticles-encapsulated drug/gene/protein for advanced controlled drug release: Current status and future perspective over emerging therapy approaches. International Journal of Biological Macromolecules. 2024:131694.

13. Bourang S, Jahanbakhsh-Godekahriz S, Asghari-Zakaria R, Parsa-Khankandi H, Noruzpour M, Calahorra J. Evaluation of antioxidant properties of essential oil, aqueous extract and metal nanoparticles biosynthesized from *F. vulgare* and their anticancer effect on two breast cancer cell lines (Sum-159, Hs-578T). Agricultural Biotechnology Journal. 2024;16(1):235–66.

14. Haider AJ, Al-Kinani MA, Al-Musawi S. Preparation and characterization of gold coated super paramagnetic iron nanoparticle using pulsed laser ablation in liquid method. Key Engineering Materials. 2021;886:77–85.

15. Mansuryar A, Bourang S, Noruzpour M, Ebrahimi HA, Amani A, Granados-Principal S, Calahorra J. The effect of Fe_3_O_4_ biosynthesized through the green synthesis of *Silybum marianum* and HA in the targeted delivery of 5-Fluorouracil to HCT116 cell line. DARU Journal of Pharmaceutical Sciences. 2025;33(2):1–18.

16. Sagar NA, Tarafdar S, Agarwal S, Tarafdar A, Sharma S. Polyamines: functions, metabolism, and role in human disease management. Medical Sciences. 2021;9(2):44.

17. Jin Y, Wang X, Kromer AP, Müller JT, Zimmermann C, Xu Z, Hartschuh A, Adams F, Merkel OM. Role of Hydrophobic Modification in Spermine-Based Poly (β-amino ester) s for siRNA Delivery and Their Spray-Dried Powders for Inhalation and Improved Storage. Biomacromolecules. 2024.

18. Zhao X, Xu Q, Wang Q, Liang X, Wang J, Jin H, Man Y, Guo D, Gao F, Tang X. Induced Self-Assembly of Vitamin E-Spermine/siRNA Nanocomplexes via Spermine/Helix Groove-Specific Interaction for Efficient siRNA Delivery and Antitumor Therapy. Advanced Healthcare Materials. 2024;13(11):2303186.

19. Bourang S, Noruzpour M, Azizi S, Yaghoubi H, Ebrahimi HA. Synthesis and *in vitro* characterization of PCL-PEG-HA/FeCo magnetic nanoparticles encapsulating curcumin and 5-FU. Nanomedicine Journal. 2024;11(2).

20. Noruzpour M, Zakaria RA, Zare N, Bourang S, Ebrahimi HA, Granados-Principal S. Delivery of *Moringa oleifera* extract via PLA-PEG-FA/Chitosan-PLA NPs into breast cancer cell lines. BioNanoScience. 2025;15(2):287.

21. Parveen S, Sahoo SK. Long circulating chitosan/PEG blended PLGA nanoparticle for tumor drug delivery. European journal of pharmacology. 2011;670(2-3):372–83.

22. Chang C-W, Choi D, Kim WJ, Yockman JW, Christensen LV, Kim Y-H, Kim SW. Non-ionic amphiphilic biodegradable PEG–PLGA–PEG copolymer enhances gene delivery efficiency in rat skeletal muscle. Journal of Controlled Release. 2007;118(2):245–53.

23. Abebe DG, Kandil R, Kraus T, Elsayed M, Merkel OM, Fujiwara T. Three-layered biodegradable micelles prepared by two-step self-assembly of PLA-PEI-PLA and PLA-PEG-PLA triblock copolymers as efficient gene delivery system. Macromolecular bioscience. 2015;15(5):698–711.

24. Fadaka A, Ajiboye B, Ojo O, Adewale O, Olayide I, Emuowhochere R. Biology of glucose metabolization in cancer cells. Journal of Oncological Sciences. 2017;3(2):45–51.

25. Komuro H, Sasano T, Horiuchi N, Yamashita K, Nagai A. The effect of glucose modification of hydroxyapatite nanoparticles on gene delivery. Journal of Biomedical Materials Research Part A. 2019;107(1):61–6.

26. Siafaka PI, Üstündağ Okur N, Karavas E, Bikiaris DN. Surface modified multifunctional and stimuli responsive nanoparticles for drug targeting: current status and uses. International journal of molecular sciences. 2016;17(9):1440.

27. Al-Nemrawi NK, Altawabeyeh RM, Darweesh RS. Preparation and characterization of docetaxel-PLGA nanoparticles coated with folic acid-chitosan conjugate for cancer treatment. Journal of Pharmaceutical Sciences. 2022;111(2):485–94.

28. Mohajeri S, Fathi Erdi A, Yaghoubi H. Targeted Gene Delivery to MCF-7 Cells via Polyspermine-PEG-Glucose/DNA Nanoparticles: Preparation and Characterization. Molecular Biotechnology. 2025:1–19.

29. Amani A, Kabiri T, Shafiee S, Hamidi A. Preparation and characterization of PLA-PEG-PLA/PEI/DNA nanoparticles for improvement of transfection efficiency and controlled release of DNA in gene delivery systems. Iranian journal of pharmaceutical research: IJPR. 2019;18(1):125.

30. Constante CK, Rodríguez J, Sonnenholzner S, Domínguez-Borbor C. Adaptation of the methyl thiazole tetrazolium (MTT) reduction assay to measure cell viability in Vibrio spp. Aquaculture. 2022;560:738568.

31. Bourang S, Asadian S, Noruzpour M, Mansuryar A, Azizi S, Ebrahimi HA, Amani Hooshyar V. PLA-HA/Fe_3_O_4_ magnetic nanoparticles loaded with curcumin: physicochemical characterization and toxicity evaluation in HCT116 colorectal cancer cells. Discover Applied Sciences. 2024;6(4):186.

32. Amani A, Dustparast M, Noruzpour M, Zakaria RA, Ebrahimi HA. Design and invitro characterization of green synthesized magnetic nanoparticles conjugated with multitargeted poly lactic acid copolymers for co-delivery of siRNA and paclitaxel. European Journal of Pharmaceutical Sciences. 2021;167:106007.

33. Bourang S, Jahanbakhsh Godehkahriz S, Noruzpour M, Asghari Zakaria R, Granados-Principal S. Anticancer properties of copolymer nanoparticles loaded with *Foeniculum vulgare* derivatives in Hs578T and SUM159 cancer cell lines. Cancer Nanotechnology. 2025;16(1):1–28.

34. Mohajeri S, Yaghoubi H, Bourang S, Noruzpour M. Multifunctional magnetic nanocapsules for dual delivery of siRNA and chemotherapy to MCF-7 cells (Breast cancer cells). Naunyn-Schmiedeberg’s Archives of Pharmacology. 2025:1–23.

35. Chong ZX, Yeap SK, Ho WY. Transfection types, methods and strategies: a technical review. PeerJ. 2021;9:e11165.

36. Antoniou J, Liu F, Majeed H, Qi J, Yokoyama W, Zhong F. Physicochemical and morphological properties of size-controlled chitosan–tripolyphosphate nanoparticles. Colloids and Surfaces A: Physicochemical and Engineering Aspects. 2015;465:137–46.

37. Danaei M, Dehghankhold M, Ataei S, Hasanzadeh Davarani F, Javanmard R, Dokhani A, Khorasani S, Mozafari M. Impact of particle size and polydispersity index on the clinical applications of lipidic nanocarrier systems. Pharmaceutics. 2018;10(2):57.

38. Emami J, Kazemi M, Hasanzadeh F, Minaiyan M, Mirian M, Lavasanifar A. Novel pH-triggered biocompatible polymeric micelles based on heparin–α-tocopherol conjugate for intracellular delivery of docetaxel in breast cancer. Pharmaceutical Development and Technology. 2020;25(4):492–509.

39. Solano-Gálvez SG, Abadi-Chiriti J, Gutiérrez-Velez L, Rodríguez-Puente E, Konstat-Korzenny E, Álvarez-Hernández D-A, Franyuti-Kelly G, Gutiérrez-Kobeh L, Vázquez-López R. Apoptosis: activation and inhibition in health and disease. Medical Sciences. 2018;6(3):54.

40. Mundel R, Thakur T, Chatterjee M. Emerging uses of PLA–PEG copolymer in cancer drug delivery. 3 Biotech. 2022;12(2):41.

41. Cavallaro G, Sardo C, Craparo EF, Porsio B, Giammona G. Polymeric nanoparticles for siRNA delivery: Production and applications. International journal of pharmaceutics. 2017;525(2):313–33.

42. Liu C, Zhang L, Zhu W, Guo R, Sun H, Chen X, Deng N. Barriers and strategies of cationic liposomes for cancer gene therapy. Molecular Therapy Methods & Clinical Development. 2020;18:751–64.

43. Casper J, Schenk SH, Parhizkar E, Detampel P, Dehshahri A, Huwyler J. Polyethylenimine (PEI) in gene therapy: Current status and clinical applications. Journal of Controlled Release. 2023;362:667–91.

44. Alnasraui AHF, Joe IH, Al-Musawi S. Design and synthesize of folate decorated Fe3O4@ Au-DEX-CP nano formulation for targeted drug delivery in colorectal cancer therapy: in vitro and in vivo studies. Journal of Drug Delivery Science and Technology. 2023;87:104798.

45. Bernkop-Schnürch A. Strategies to overcome the polycation dilemma in drug delivery. Advanced drug delivery reviews. 2018;136:62–72.

46. Ucal S. Polyamine analogues as anticancer agents: Itä-Suomen yliopisto; 2017.

47. Abdulwahid FS, Haider AJ, Al-Musawi S. Effect of laser parameter on Fe3O4 NPs formation by pulsed laser ablation in liquid. 2023.

48. Wadhawan A, Singh J, Sharma H, Handa S, Singh G, Kumar R, Barnwal RP, Pal Kaur I, Chatterjee M. Anticancer biosurfactant-loaded PLA–PEG nanoparticles induce apoptosis in human MDA-MB-231 breast cancer cells. ACS omega. 2022;7(6):5231–41.

49. Ramalho MJ, Alves B, Andrade S, Lima J, Loureiro JA, Pereira MC. Folic-acid-conjugated poly (lactic-co-glycolic acid) nanoparticles loaded with gallic acid induce glioblastoma cell death by reactive-oxygen-species-induced stress. Polymers. 2024;16(15):2161.

50. Jain AK, Thareja S. *In vitro* and *in vivo* characterization of pharmaceutical nanocarriers used for drug delivery. Artificial cells, nanomedicine, and biotechnology. 2019;47(1):524–39.

51. Rostami N, Gomari MM, Abdouss M, Moeinzadeh A, Choupani E, Davarnejad R, Heidari R, Bencherif SA. Synthesis and characterization of folic acid-functionalized DPLA-co-PEG nanomicelles for the targeted delivery of letrozole. ACS Applied Bio Materials. 2023;6(5):1806–15.

52. Le-Vinh B, Le N-MN, Nazir I, Matuszczak B, Bernkop-Schnürch A. Chitosan based micelle with zeta potential changing property for effective mucosal drug delivery. International journal of biological macromolecules. 2019;133:647–55.

53. Okami K, Fumoto S, Yamashita M, Nakashima M, Miyamoto H, Kawakami S, Nishida K. One-Step Formation Method of Plasmid DNA-Loaded, Extracellular Vesicles-Mimicking Lipid Nanoparticles Based on Nucleic Acids Dilution-Induced Assembly. Cells. 2024;13(14):1183.

54. Alshamsan A, Haddadi A, Hamdy S, Samuel J, El-Kadi AO, Uludag H, Lavasanifar A. STAT3 silencing in dendritic cells by siRNA polyplexes encapsulated in PLGA nanoparticles for the modulation of anticancer immune response. Molecular pharmaceutics. 2010;7(5):1643–54.

55. Lin PH, Huang C, Hu Y, Ramanujam VS, Lee E-S, Singh R, Milbreta U, Cheung C, Ying JY, Chew SY. Neural cell membrane-coated DNA nanogels as a potential target-specific drug delivery tool for the central nervous system. Biomaterials. 2023;302:122325.

56. Alipournazari P, Pourmadadi M, Abdouss M, Rahdar A, Pandey S. Enhanced delivery of doxorubicin for breast cancer treatment using pH-sensitive starch/PVA/g-C3N4 hydrogel. International Journal of Biological Macromolecules. 2024;265:130901.

57. Kesharwani P, Halwai K, Jha SK, Al Mughram MH, Almujri SS, Almalki WH, Sahebkar A. Folate-engineered chitosan nanoparticles: next-generation anticancer nanocarriers. Molecular Cancer. 2024;23(1):244.

58. Soltani M, Ahmadzadeh N, Nasiraei Haghighi H, Khatamian N, Homayouni Tabrizi M. Targeted cancer therapy potential of quercetin-conjugated with folic acid-modified nanocrystalline cellulose nanoparticles: a study on AGS and A2780 cell lines. BMC biotechnology. 2025;25(1):29.

59. Schmidts T, Dobler D, von den Hoff S, Schlupp P, Garn H, Runkel F. Protective effect of drug delivery systems against the enzymatic degradation of dermally applied DNAzyme. International journal of pharmaceutics. 2011;410(1-2):75–82.

60. Kondinskaia DA, Gurtovenko AA. Supramolecular complexes of DNA with cationic polymers: The effect of polymer concentration. Polymer. 2018;142:277–84.

61. Chandrasekaran AR, Halvorsen K. Nuclease degradation analysis of DNA nanostructures using gel electrophoresis. Current protocols in nucleic acid chemistry. 2020;82(1):e115.

